# A Comprehensive Benchmarking Study on Computational Tools for Cross-omics Label Transfer from Single-cell RNA to ATAC Data

**DOI:** 10.1101/2024.02.01.578507

**Authors:** Yuge Wang, Hongyu Zhao

## Abstract

With continuous progress of single-cell chromatin accessibility profiling techniques, scATAC-seq has become more commonly used in investigating regulatory genomic regions and their involvement in developmental, evolutionary, and disease-related processes. At the same time, accurate cell type annotation plays a crucial role in comprehending the cellular makeup of complex tissues and uncovering novel cell types. Unfortunately, the majority of existing methods primarily focus on label transfer within scRNA-seq datasets and only a limited number of approaches have been specifically developed for transferring labels from scRNA-seq to scATAC-seq data. Moreover, many methods have been published for the joint embedding of data from the two modalities, which can be used for label transfer by adding a classifier trained on the latent space. Given these available methods, this study presents a comprehensive benchmarking study evaluating 27 computational tools for scATAC-seq label annotations through tasks involving single-cell RNA and ATAC data from various human and mouse tissues. We found that when high quality paired data were available to transfer labels across unpaired data, Bridge and GLUE were the best performers; otherwise, bindSC and GLUE achieved the highest prediction accuracy overall. All these methods were able to use peak-level information instead of purely relying on the gene activities from scATAC-seq. Furthermore, we found that data imbalance, cross-omics dissimilarity on common cell types, data binarization, and the introduction of semi-supervised strategy usually had negative impacts on model performance. In terms of scalability, we found that the most time and memory efficient methods were Bridge and deep-learning-based algorithms like GLUE. Based on the results of this study, we provide several suggestions for future methodology development.

## 1 Introduction

The integration of single-cell RNA sequencing (scRNA-seq) and single-cell assay for transposase-accessible chromatin sequencing (scATAC-seq) has become instrumental in unraveling the intricacies of cellular heterogeneity and regulatory mechanisms at a single-cell resolution [1-5]. However, the accurate annotation of scATAC-seq cells remains a challenge and involves a combination of automated annotations from computational tools and subsequent manual corrections [6]. While numerous tools exist for annotating cell types in scRNA-seq data [7, 8], the options tailored specifically for scATAC-seq data are limited. In addition, many methods have been designed for computational integration of scRNA-seq and scATAC-seq data, typically resulting in a joint embedding of cells from both modalities in a low-dimensional space [9-11]. Once this joint embedding is obtained, label transfer can be easily conducted such as through an external k nearest neighbor (kNN) classifier trained on the latent space. As scATAC-seq matures and becomes widely adopted in single-cell studies, it is crucial to conduct a comprehensive evaluation of the performance of methods from both categories in annotating scATAC-seq data.

Within the first category, there are two different types of annotation tools applicable for the annotation of scATAC-seq data. The first type comprises tools initially designed for scRNA-seq data (intra-modality annotation), while the second type includes tools specifically designed for scATAC-seq data (cross-modality annotation). Popular methods in the first type are Seurat v3 [12], Conos [13] and scGCN [14] and two representative methods in the second type include scJoint [15] and Bridge [16]. In contrast to other methods that directly transfer labels from scRNA-seq to scATAC-seq after consolidating the feature set through gene activity calculation, Bridge utilizes multimodal data as a bridge. This approach helps circumvent potential information loss and inaccuracies in assumptions about feature relationships between the two modalities.

For the second category, there are also two types of tools that can perform joint embedding of scRNA-seq and scATAC-seq data. For tools belonging to the first type, they were originally designed for integrating different scRNA-seq data with batch effects, such as LIGER [17], scVI [18], SCALEX [19] and scDML [20]. These methods can be applied to integrate scRNA-seq and scATAC-seq data once the ATAC data are transformed to gene activities. The second type includes tools that were published more recently and designed for integrating multimodal single-cell data. Although methods under this type advertised themselves for the ability to integrate scRNA-seq and scATAC-seq data, some of them still require gene activity calculation for the ATAC data in order to directly align the feature sets of two modalities, such as cross-modal AE [21], uniPort [22] and scMC [23]. There are also other methods that can utilize the ATAC peak data such as MMD-MA [24], UnionCom [25], SCOT [26, 27], Pamona [28], MultiMAP [29], scDART [30], GLUE [31], bindSC [32], UINMF [33], MultiVI [34], Cobolt [35] and StabMap [36], among which MultiVI and Cobolt work like Bridge that require an additional paired data to integrate unpaired RNA and ATAC data. GLUE and StabMap have two versions of themselves. One version does not require paired data and the other version can use paired data to help better integrate unpaired data if available.

Previously, we published a benchmarking study on this topic on a smaller scale, using only five datasets and focusing solely on the five methods from the first category[37]. This current study represents a substantial expansion, encompassing all the 27 computational tools mentioned above. We employed a more extensive selection of single-cell RNA and ATAC data from both human and mouse tissues, each with available cell type annotations serving as the ground truth. The data we collected included nine paired data (multimodal) where scATAC-seq and scRNA-seq were simultaneously measured in each single cell and 18 unpaired data (unimodal) where scATAC-seq and scRNA-seq were separately measured from the same tissue. Among the unpaired samples, nine had corresponding paired data from the same species and tissue, facilitating the evaluation of methods that require additional paired data for direct label transfer from RNA to ATAC or integration of data from the two modalities. We evaluated the performance of different methods on both annotation accuracy and scalability. For accuracy, we considered both accuracy metrics calculated on common cell types and metrics calculated on ATAC-specific cell types. For scalability, we compared running time and peak memory usage across a wide range of data sizes.

Beyond benchmarking and ranking, we performed additional comparisons and designed experiments to understand how model performance is affected by different data properties and method properties. For data properties, we considered three aspects, namely the proportion of modality-specific cell types, the discrepancy in the common cell type compositions between modalities, and the cross-omics dissimilarity of common cell types. For method properties, we discussed how the incorporation of paired data, peak level information of ATAC data, and the introduction of semi-supervised training would affect label transfer accuracy. At the end of this paper, we extensively discuss observations related to both method and data properties and provide guidance for users in selecting or developing suitable tools for cell type annotation in scATAC-seq data.

## 2 Results

### 2.1 Cross-omics label transfer benchmarking design

Here, we provide a comprehensive and quantitative evaluation of 27 computational tools for cross-omics label transfer from scRNA-seq data to scATAC-seq data. The overview of our benchmarking design is shown in Figure 1.

**Figure 1.**
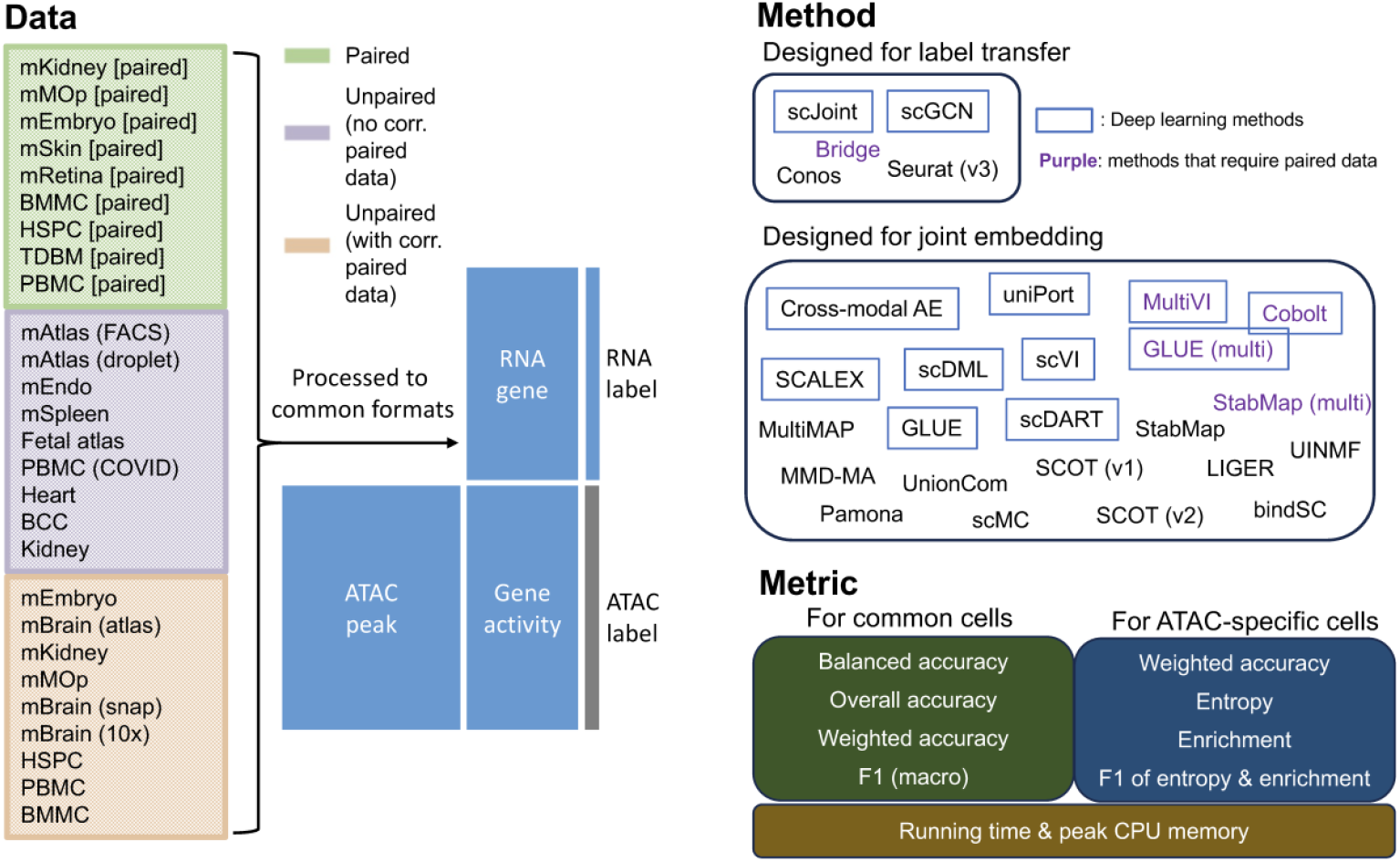
Overview of the benchmarking design in terms of data, methods, and metrics. In this study, we evaluated 27 methods, where five methods are designed for label transfer which means that the predicted labels are included in the outputs of these algorithms, and the other 22 methods are designed for joint embedding of scRNA-seq and scATAC-seq data. To perform label transfer, we first obtained the joint embedding and then used that to train a kNN classifier (k=30). Among both categories of methods, there are in total five methods that involve additional paired data, so they were only run on the third group of datasets, where unpaired RNA and ATAC data had corresponding paired data. The full details about the 27 methods can be found in Table 1 and Methods (4.2).

We collected a variety of scRNA-seq and scATAC-seq datasets from both human and mouse tissues, including brain [38-40], primary motor cortex [41], retina [32], skin [42], kidney [43-45], spleen [29, 46], blood [47, 48], bone marrow [3, 49-51], arteries [52], heart [53], embryo [54-56], basal cell carcinoma [57, 58]. Apart from tissue-specific data, we also collected a mouse atlas [59, 60] and a human fetal atlas dataset [61, 62]. All data can be divided into three groups: paired data, unpaired data with no corresponding paired data from the same tissue, and unpaired data with corresponding paired data from the same tissue. For the paired data, we did not use the pairing information when applying each method. For the unpaired data with corresponding paired data, the pairing information of the paired data was used for methods that can integrate both unpaired and paired data. Before running all methods, we processed all data to common formats, which are RNA gene expression count matrix, ATAC peak count matrix, ATAC gene activity matrix, RNA cell type labels, and ATAC cell type labels. The ATAC cell type labels were only used for evaluation. The full details about the data we collected and the steps we used to process the data can be found in Supplementary Table S1 and Methods (4.1).

**Table 1.**
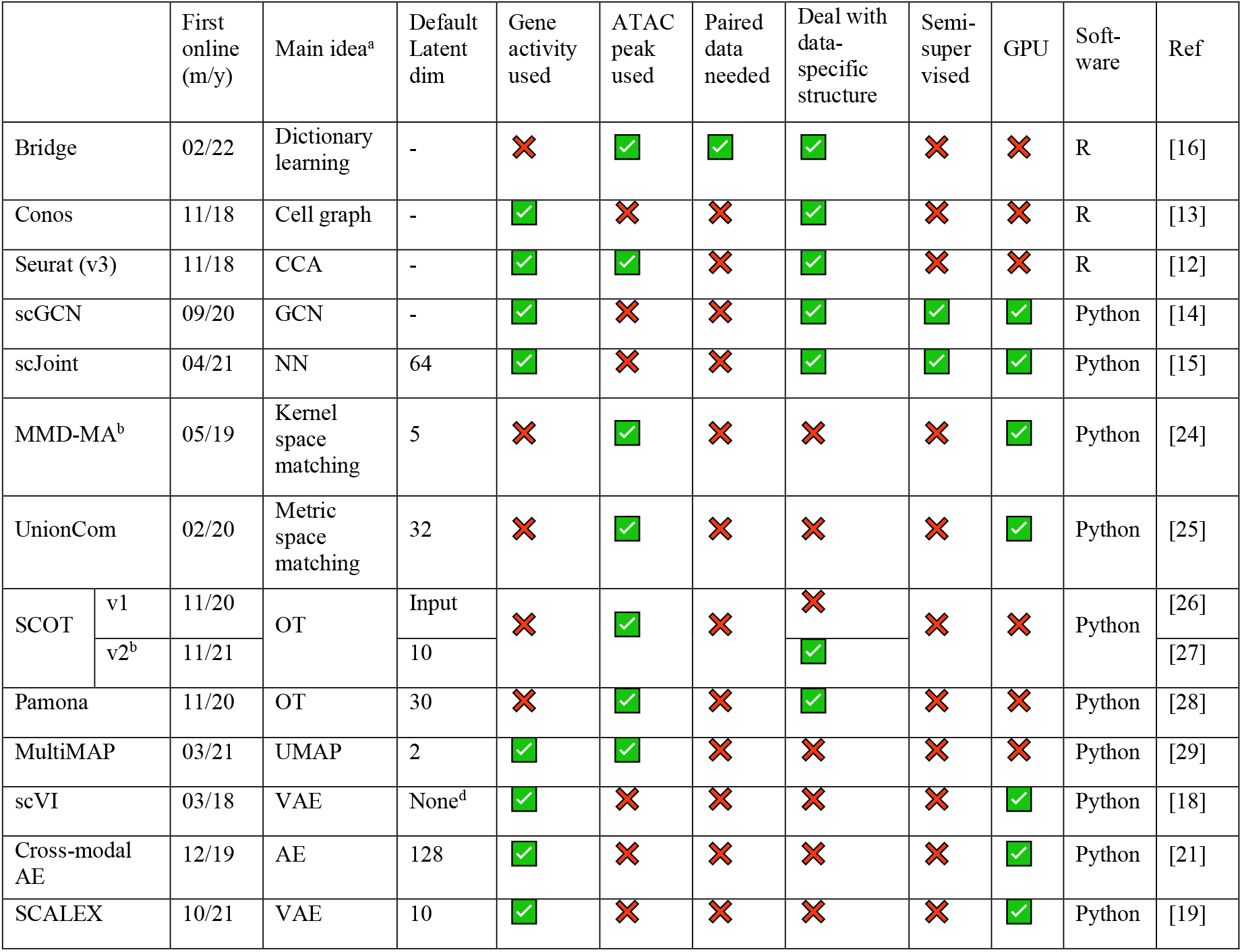
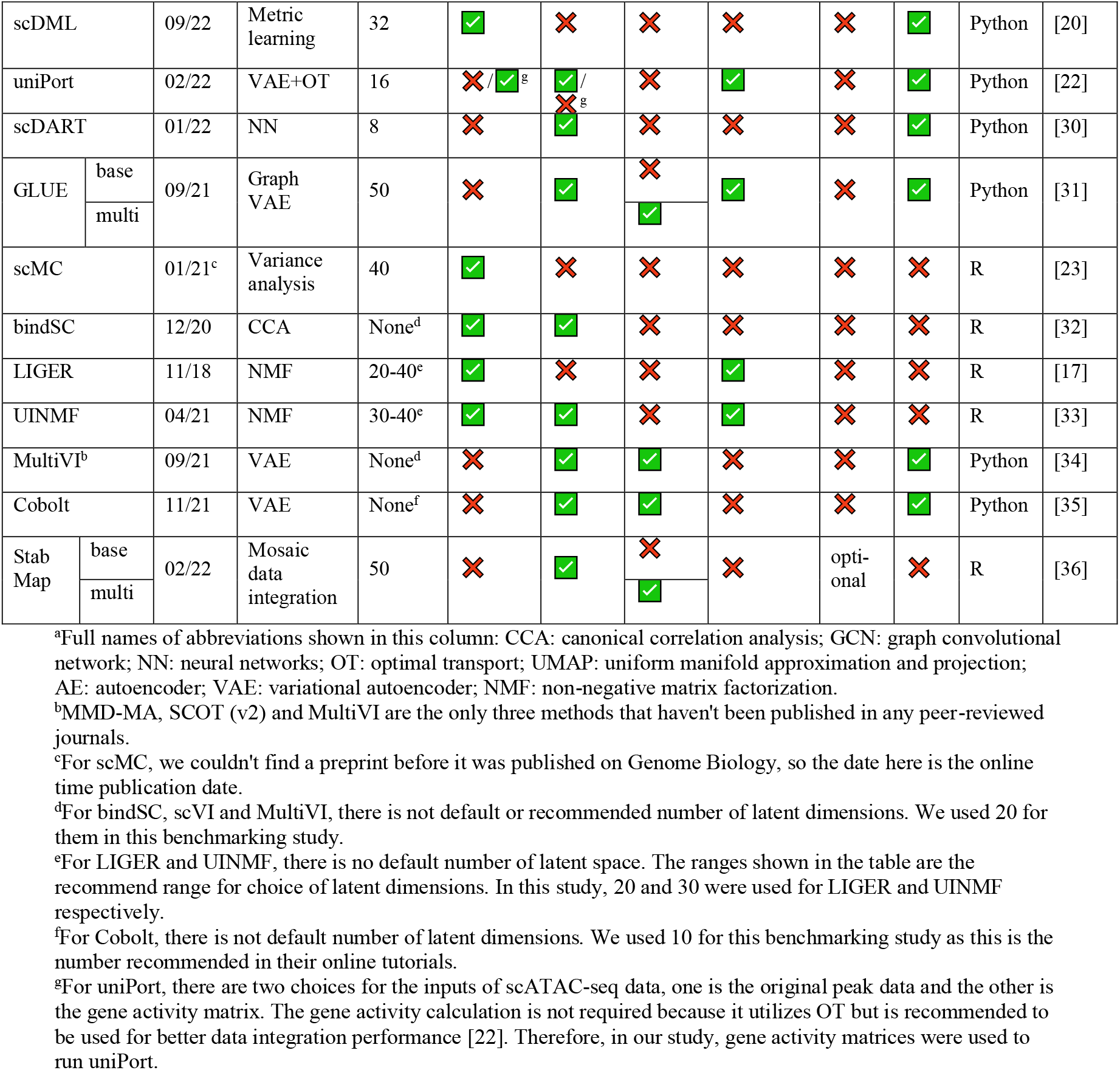
Overview of methods considered in this benchmarking study.

In addition to the 27 methods, we used kNN and random classifiers as the baseline competitors. For kNN classifiers, all common features between scRNA-seq and gene activity matrix calculated from the scATAC-seq data were used for training. For random classifiers, labels were predicted based on the background probabilities of cell types in the scRNA-seq data.

After getting the predicted probability matrix from each method for ATAC cells, we assessed the prediction accuracy for common cells and ATAC-specific cells separately. For common cells, we calculated four metrics, namely balanced accuracy, overall accuracy, weighted accuracy and F1 (macro) of precision and recall. For the first, second and fourth metrics, they were calculated based on the predicted label of each cell, which was the cell type whose predicted probability was the highest. For weighted accuracy, we considered the similarity among cell types by calculating the weighted average of the entire predicted probability vector of each cell. Therefore, even though a predicted label was false, the score could be high if similar cell types had higher predicted probabilities. For ATAC-specific cells, we calculated four metrics as well, namely weighted accuracy, entropy score, enrichment score and F1 of entropy and enrichment. In addition to accuracy evaluation, we measured the scalability of each method through the running time and peak CPU memory usage across selected gradients of sample sizes. Details about the metrics we used can be found in Methods (4.3).

In addition to assessing the overall performance of the 27 methods across the 27 datasets (tasks), we conducted an in-depth analysis to explore factors that could have played important roles in cross-modality label transfer from the aspects of both data properties and model properties.

### 2.2 Performance across different tissues

To equally weigh the contribution of different metrics and tasks to the performance comparison among different methods, we min-max scaled the values of each metric within a task to make sure that the scaled metric covered the entire range from 0 to 1 for a task. The overall prediction accuracy on common cell types for each task and method combination was calculated by averaging across all the four min-max scaled metrics. Results for each of the three groups of tasks (datasets) mentioned previously are shown in Figure 2. For the original unscaled metrics, they can be found in Supplementary Figure S2-S4.

**Figure 2.**
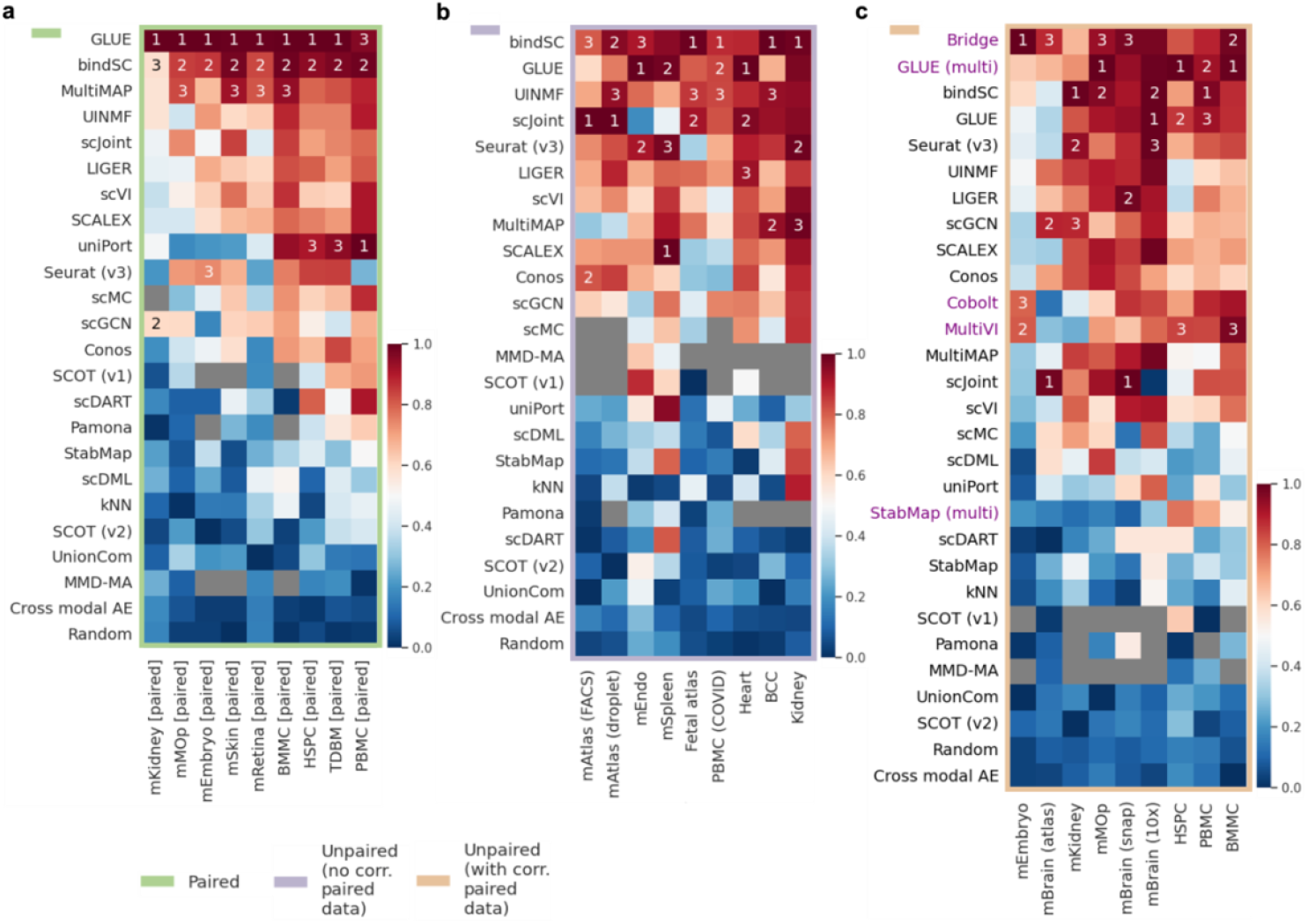
Performance of 27 methods on cross-omics label transfer (RNA to ATAC) across 27 mouse and human datasets. Datasets were divided into three groups: paired datasets (pairing information not used), unpaired datasets without corresponding paired data available, and unpaired datasets with corresponding paired data available. ‘Corresponding’ means from the same tissue and species. (a) The average of four min-max scaled metrics evaluated on common cells (balanced accuracy, overall accuracy, weighted accuracy and F1 (macro)) for paired datasets. (b) The average of four min-max scaled metrics evaluated on common cells for unpaired datasets with no corresponding paired data. (c) The average of four min-max scaled metrics evaluated on common cells for unpaired datasets with corresponding paired data. For each heatmap, methods (rows) are placed from top to bottom in the descending order of their averaged values across all datasets (columns) in that heatmap and methods ranked top three in each dataset are marked by Arabic numbers. Heatmaps for each metric in their original scales, including those for common cells and for ATAC-specific cell, can be found in Supplementary Figure S2-S4.

For paired data (Figure 3a), GLUE, bindSC and MultiMAP performed the best, followed by UINMF and scJoint. GLUE and bindSC were the only two methods that consistently showed up in the top three performers across all tasks. Especially for GLUE, it was the best performer in eight out of nine tasks. The performance of uniPort seemed to vary depend on the specific task. Among the nine tasks, it was among the top three performers in three of them.

**Figure 3.**
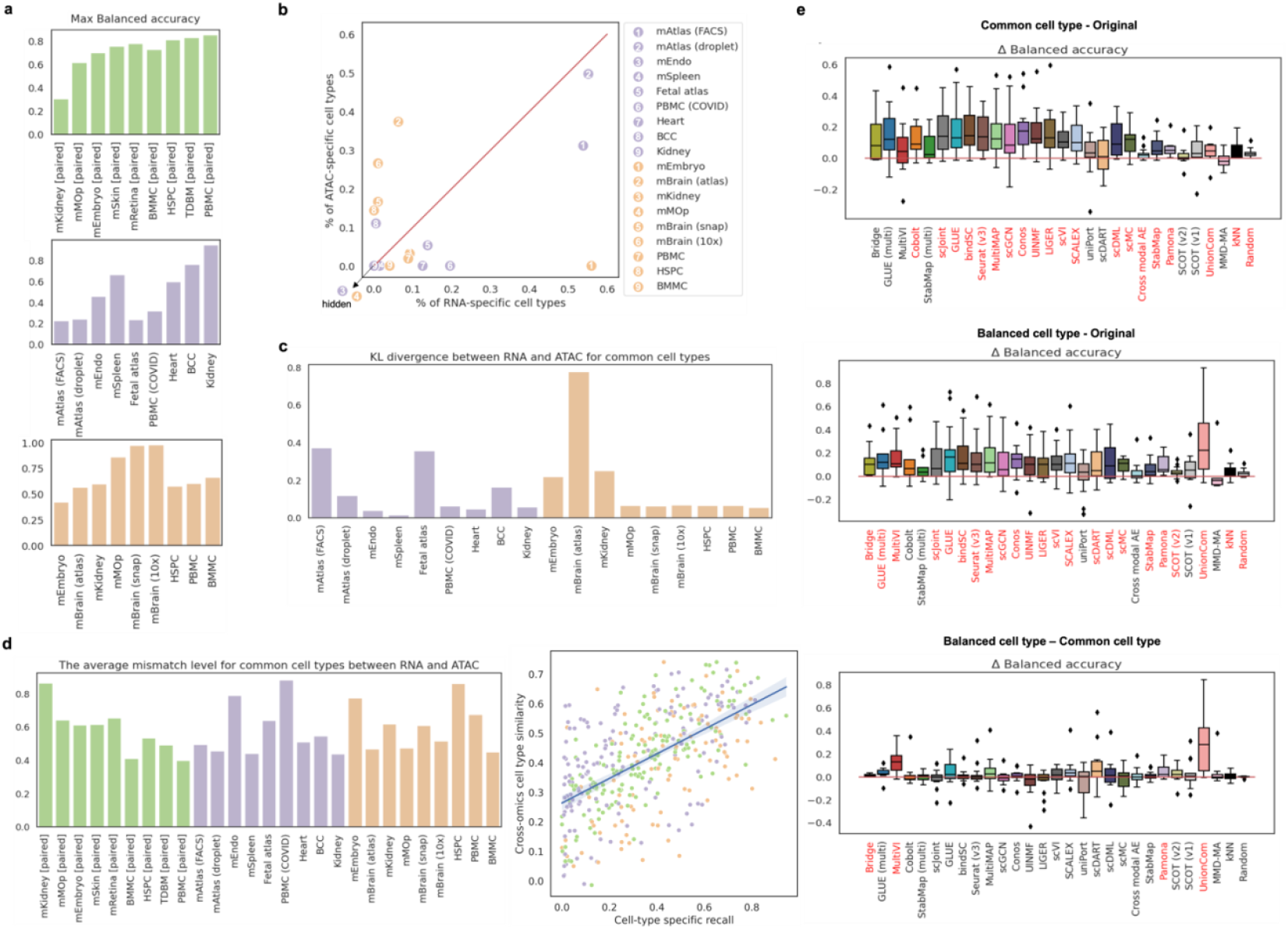
The impact of data properties, including data imbalance and cross-omics cell type similarity, on prediction accuracy. (a) The maximum values of balanced accuracy achieved on each dataset across all the methods. Information of other metrics can be found in Supplementary Figure S1. (b) The proportions of modality-specific cell types for unpaired datasets. (c) The discrepancy in cell type compositions between two modalities for unpaired datasets measured by KL divergence. (d) The influence of cross-omics mismatch level on prediction accuracy. On the left is the cross-omics mismatch level, which is defined as 1 minus the cross-omics similarity, averaged across all common cell types between RNA and ATAC for each task. On the right is the relationship between cross-omics cell type similarity and cell-type specific recall. Each dot is the average recall across all methods for a specific common cell type in one dataset. The cross-omics similarity for each common cell type was calculated by the Pearson correlation using 2,000 common HVGs. (e) The impact of manually removing cell type imbalance on balanced accuracy. ‘Common cell type’ means modality-specific cell types were removed and ‘balanced cell type’ means cell type compositions were made the same between two modalities. Pairwise comparisons among these two cases and the original case for each method and dataset combination were performed. For all boxplots, methods marked in red along the x-axes are methods that achieved significant positive differences (p-value threshold: 0.05; statistical test: one-sample t-test). The impact on other metrics can be found in Supplementary Figure S6-S8.

For unpaired data without corresponding paired data available (Figure 3b), the top three performers were bindSC, GLUE and UINMF, with scJoint and Seurat (v3) following closely behind when considering the metrics calculated on common cells. In terms of F1 of entropy and enrichment calculated on ATAC-specific cells (Supplementary Figure S3b), random classifiers undoubtedly achieved the highest scores, followed by scGCN and uniPort, indicating that these two methods performed most similar to a random classifier. Moreover, we observed that scJoint had very low F1 of entropy and enrichment and relatively high weighted accuracy on ATAC-specific cells. This suggests that scJoint tended to classify ATAC-specific cell types to existing cell types that were biologically similar to them.

For unpaired data with corresponding paired data available (Figure 3c), we were able to assess all the 27 methods including the five methods that required paired data as a ‘bridge’ to transfer information across omics. For metrics calculated using common cells, the top three performers were Bridge, GLUE (multi) and bindSC, followed by GLUE and Seurat (v3). Regarding metrics computed with ATAC-specific cells (Supplementary Figure S4b), methods that performed most akin to a random classifier were scGCN and scDML. As for scJoint, similar to what was observed in the second type of datasets, it achieved relatively low F1 of entropy and enrichment and high weighted accuracy on ATAC-specific cells.

Note that for LIGER, there were two options when selecting highly variable genes (HVGs), which was one of the steps of the algorithm used in LIGER. One option was selecting using both the RNA and ATAC data and the other was only using the RNA data. We found there were no significant differences between these two versions of LIGER across all metrics (Supplementary Figure S5a), so we used the first version of LIGER throughout the entire paper. Like LIGER, StabMap also had two options when selecting the reference space to map onto. We found that when using ATAC as the reference, seven out eight metrics achieved higher values than using RNA as the reference (Supplementary Figure S5b). Therefore, we used ATAC as the reference space for StabMap in our study.

### 2.3 The impact of data properties

Previously, we min-max scaled each metric to equally weigh the contributions of different metrics to the overall prediction accuracy. However, if we take a look at the metrics in their original scales, we will find for some tasks, even the best performers had a very low prediction accuracy (Figure 3a and Supplementary Figure S1). For example, for mKidney [paired], mAtlas (FACS), mAtlas (droplet), fetal atlas and PBMC (COVID), the maximum balanced accuracy was only around 0.2 to 0.3. We found this was due to the properties of the datasets used in the task, including the level of data imbalance and cross-omics cell type dissimilarity (Figure 3).

#### 2.3.1 Data imbalance

There were two levels of data imbalance: the proportion of modality-specific cell types (Figure 3b) and the discrepancy in the compositions of common cell types (Figure 3c). We found that datasets with either high proportions of modality-specific cell types or substantial discrepancies in common cell type compositions exhibited poor performance across all methods. For example, mAtlas (FACS), mAtalas (droplet) and mEmbryo contained over 50% of RNA-specific cell types. The discrepancy in common cell type compositions of fetal atlas and mBrain (atlas) ranked the highest among all human datasets and all mouse datasets, respectively.

In addition to the aforementioned empirical evidence, we studied the impact of cell type imbalance on prediction accuracy by manually adjusting data imbalance at the two levels. First, we equalized the sets of cell types between the two modalities by removing modality-specific cell types from both training and evaluation, denoting this as the ‘common cell type’ experiment. Second, we further manually selected cells to ensure that RNA and ATAC data had the same number of cells for each common cell type and referred to as the ‘balanced cell type’ experiment.

When comparing the ‘common cell type’ with the original data (Figure 3e top and Supplementary Figure S6), we observed that 18 out of the 27 methods (66.7%) demonstrated increased prediction accuracy for at least one metric assessed on common cells. This suggests that merely removing modality-specific cell types could significantly improve the prediction accuracy for common cells. In the comparison of the ‘balanced cell type’ with the original data (Figure 3e middle and Supplementary Figure S7), we noted an increase in the number of methods that had improved prediction accuracy (22 out of 27, 81.5%). The methods whose performance remained unaffected in both experiments were uniPort, SCOT (v1) and MMD-MA. Moreover, we investigated whether balancing cell type compositions could improve the prediction accuracy beyond the removal of modality-specific cell types by directly comparing the results of the ‘balanced cell type’ with the results of the ‘common cell type’ experiment (Figure 3e bottom and Supplementary Figure S8). This time, fewer methods showed significant improvement in at least one metric (14 out of 27, 51.9%). Especially for balanced accuracy, significant changes were only observed for Bridge, MultiVI, Pamona and UnionCom, with changes being non-negligible for only MultiVI and UnionCom.

#### 2.3.2 Cross-omics cell type dissimilarity

Although some datasets didn’t exhibit high data imbalance, their cross-omics cell type dissimilarity was very high, contributing to their poor label transfer performance across all the methods. For example, mEndo and PBMC (COVID) were the tasks that had the highest mismatch scores for common cell types between RNA and ATAC among all unpaired mouse and human tasks, respectively. So as for mKidney [paired] in the paired data group. Overall, we found that the cross-omics similarity for common cell types was positively related with the prediction accuracy (Figure 3d right, correlation: 0.59, p-value: 6.7e-40).

We noted that different methods processed single-cell ATAC data differently. For example, scDART, MultiVI and StabMap binarized the ATAC peak matrix and scJoint even binarized both the RNA and ATAC gene matrices. According to scJoint, data binarization of both RNA and ATAC improved its performance because the distributions of two modalities were made more similar. To investigate whether this also applied to other methods, we performed data binarization before running each method. From Supplementary Figure S9, we first observed that data binarization had minimal effect on scJoint as expected. Then, we found none of the methods benefitted from data binarization in terms of the prediction accuracy on common cells and 14 methods out of 27 (51.9%) had significantly decreased accuracy in at least one metric. The methods that were affected most by data binarization were UINMF and LIGER. Interestingly, for the metrics calculated on ATAC-specific cells, we found the weighted accuracy was only significantly and negatively affected for Bridge and the F1 of entropy and enrichment was positively affected for 13 methods, among which 10 methods’ prediction accuracy on common cells were negatively affected. The methods that were not affected by data binarization at all were GLUE (multi), MultiVI, Cobolt, scJoint, MultiMAP, scDART, scDML, SCOT (v1) and UnionCom.

### 2.4 The impact of method properties

#### 2.4.1 The involvement of paired data

Since we found both of the top two performers for common cells’ prediction accuracy in Figure 2c were methods that involved paired data as a ‘bridge’, we wanted to study how using additional paired data could affect the label transfer accuracy by directly comparing four pairs of methods: MultiVI and scVI, GLUE (multi) and GLUE, Bridge and Seurat, and StabMap (multi) and StabMap (Figure 4a and Supplementary Figure S10). We found the prediction accuracy on common cells was significantly improved for GLUE (multi) and Bridge, while the metrics for ATAC-specific cells didn’t change much. For MultiVI and StabMap (multi), their performance didn’t change significantly compared to their base versions, respectively.

**Figure 4.**
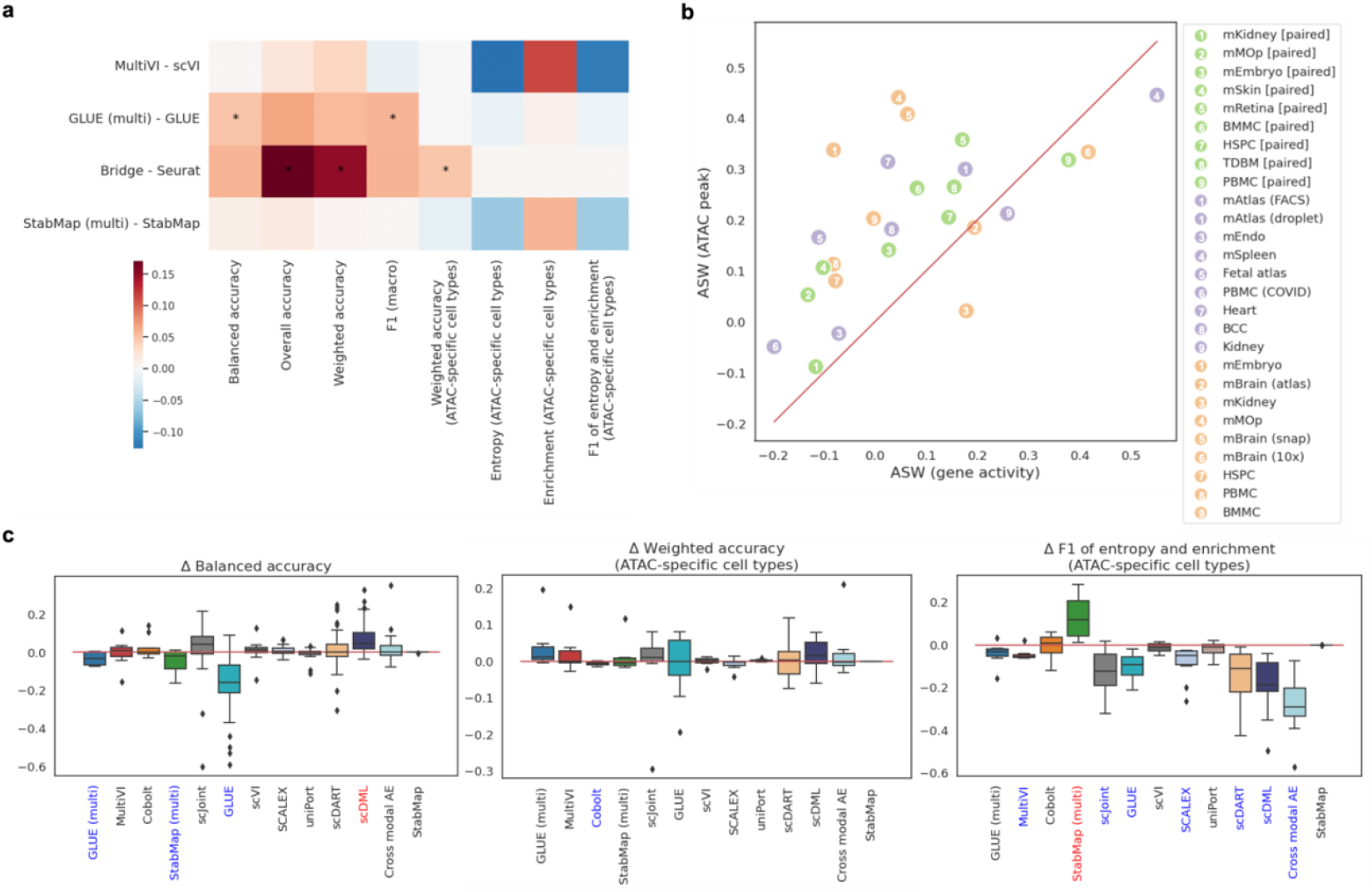
The impact of method properties, including the usage of paired data, the usage of original ATAC peak data, and the introduction of semi-supervised strategy. (a) The impact of using paired data as a ‘bridge’. The heatmap shows the differences in accuracy metrics (columns) averaged across all tissues with asterisks indicating statistical significance. (b) The comparison of average Silhouette width (ASW) between UMAPs drawn using ATAC peak data and gene activity (GA) matrix. Note that we have one less ATAC data than the total number of datasets because mAtlas (FACS) and mAtlas (droplet) shared the same ATAC data. (c) The impact of introducing semi-supervised strategy to 13 selected methods on prediction accuracy. The impact was evaluated by the differences between metrics calculated using results after and before employing the semi-supervised strategy. From left to right, the three boxplots show the differences in balanced accuracy assessed on all common cells, weighted accuracy and F1 of entropy and enrichment accuracy assessed on ATAC-specific cells. Methods marked in red and blue along the x-axes are methods that achieved positive and negative significant differences respectively. All statistical tests conducted here were one-sample t-tests with p-value threshold being 0.05.

#### 2.4.2 The utilization of the peak information

All the top performers mentioned in Section 2.2 were methods that utilized the ATAC peak-level information in some way. This is likely because the peak-level data contained more biological information than the gene-level data for scATAC-seq. Specifically, we found different cell types had better separations in UMAPs derived from the peak data compared to that from the gene activities (Supplementary Figure S11) and the ASW calculated using the peak data was significantly higher than that calculated using the gene activity data, with p-value of paired t-test being 1.12e-4 (Figure 4b and Supplementary Figure S12).

#### 2.4.3 The introduction of semi-supervised training

For cross-omics label transfer, the RNA cell type labels are assumed to be part of the known information. scGCN and scJoint are the two methods among all the five methods that are designed for label transfer that are able to utilize this information through a semi-supervised training strategy. There have also been studies arguing that using available cell labels could achieve better cell embeddings [63-65]. Therefore, we investigated whether introducing semi-supervised strategy to some of the joint embedding methods could increase the prediction accuracy.

From Figure 4c and Supplementary Figure S13, we observed that for metrics assessed on common cells, scDML was the only method that benefitted largely and significantly from semi-supervised training, while for methods including GLUE (multi), StabMap (multi) and GLUE, their prediction accuracy decreased significantly after using semi-supervised training. The weighted accuracy on ATAC-specific cells didn’t change much on any of the methods. For F1 of entropy and enrichment calculated using ATAC-specific cells, seven methods (MultiVI, scJoint, GLUE, SCALEX, scDART, scDML and cross-modal AE) had decreased values and StabMap was the only method that had increased values. Methods that were insensitive to the introduction of semi-supervised training were Cobolt, scVI, uniPort and StabMap.

### 2.5 Time and memory usage comparison

We used mMOp to assess the scalability of all the 27 methods, because this was the only task that originally contained more than 50k cells for both the RNA and the ATAC data and had corresponding paired data. Specifically, we ran each method on a series of data sizes, including 1k, 3k, 5k, 10k, 15k, 20k, and 50k cells, and recorded the running time and peak CPU memory usage.

As shown in Figure 5a, the top five methods that increased fastest in running time as the data size increased were SCOT (v2), UnionCom, Pamona, SCOT (v1) and scDART. The first four methods were designed based on optimal transport or metric space matching that involved time-consuming matrix optimization especially when data scale went up. scDART is a deep-learning-based method, but it requires the calculation of pairwise distances between cells in the latent space. The top five methods that scaled slowest as the data size increased were uniPort, SCALEX, GLUE, GLUE (multi) and Cobolt. All of them are deep-learning-based methods, but unlike scDART, none of them involved the computation of loss terms that scaled non-linearly with the data size. The first four methods even employed early stopping and iteration-based training instead of epoch-based training to control the running time (see comparisons among deep-learning-based methods in Table 2).

**Table 2.** A summary of common hyperparameters of deep-learning-based methods.

|  | Default/used epochs | Default/used batch size | Early stopping? | Optimizer | Learning rate |
| --- | --- | --- | --- | --- | --- |
| scGCN | 200 | N<sup>a</sup> | ✓ | Adam | 0.01 |
| scJoint | 20<sup>b</sup> | 256<sup>b</sup> | ✗ | SGD | 0.01<sup>b</sup> |
| scDML | 50 | 64 | ✗<sup>c</sup> | Adam | 0.01 |
| Cross-modal AE<sup>d</sup> | 100 | 128 | ✗ | SGD | 0.005 |
| GLUE | max(48/0.002/(N/batch size), 48)<sup>e</sup> | 128 | ✓ | RMSprop | 0.002 |
| scDART | 500<sup>f</sup> | 128 | ✗ | Adam | 0.0003 |
| SCALEX | 30000/(N/batch size)<sup>e</sup> | 64 | ✓ | Adam | 0.0002 |
| uniPort | 30000/(N/batch size)<sup>e</sup> | 256 | ✓ | Adam | 0.0002 |
| scVI/MultiVI | min(20000/N*400, 400) | 128 | ✗<sup>c</sup> | Adam | 0.001 |
| Cobolt | 100 | 128 | ✗ | Adam | 0.001<sup>f</sup> |

<sup>a</sup>For the entire table, we use N to indicate the total number of cells.
<sup>b</sup>For scJoint, there were no default numbers of epochs, batch size or learning rate provided. The values shown in the table were what we used as default in this study, which were chosen by referring to the values used in its real data applications. Note that for the learning rate, we used 0.001 for mMOP (both stage 1 and 3), HSPC [paired] and mEmbryo [paired] (only stage 3) since 0.01 would cause the training of scJoint to diverge on these tissues.
<sup>c</sup>By default early stopping was not used for scDML, scVI and MultiVI, but users can switch that on if desired.
<sup>d</sup>For cross-model AE, there were no code provided for this algorithm from the original authors. To implement it, we downloaded the version provided by a comment paper [9].
<sup>e</sup>For GLUE, SCALEX and uniPort, instead of fixing the number of epochs, they set the maximum number of iterations, which is defined as training one batch size samples.
<sup>f</sup>Although the default number of epochs is 700 as written in its training function, we used 500 because this was the number used in the original publication and its online tutorial.
<sup>g</sup>For Cobolt, we adjusted the learning rate down to 0.0002 for mEmbryo in all experiments and for mBrain (snap) and mKidney in the data binarization experiment because we found otherwise the training would diverge.

**Figure 5.**
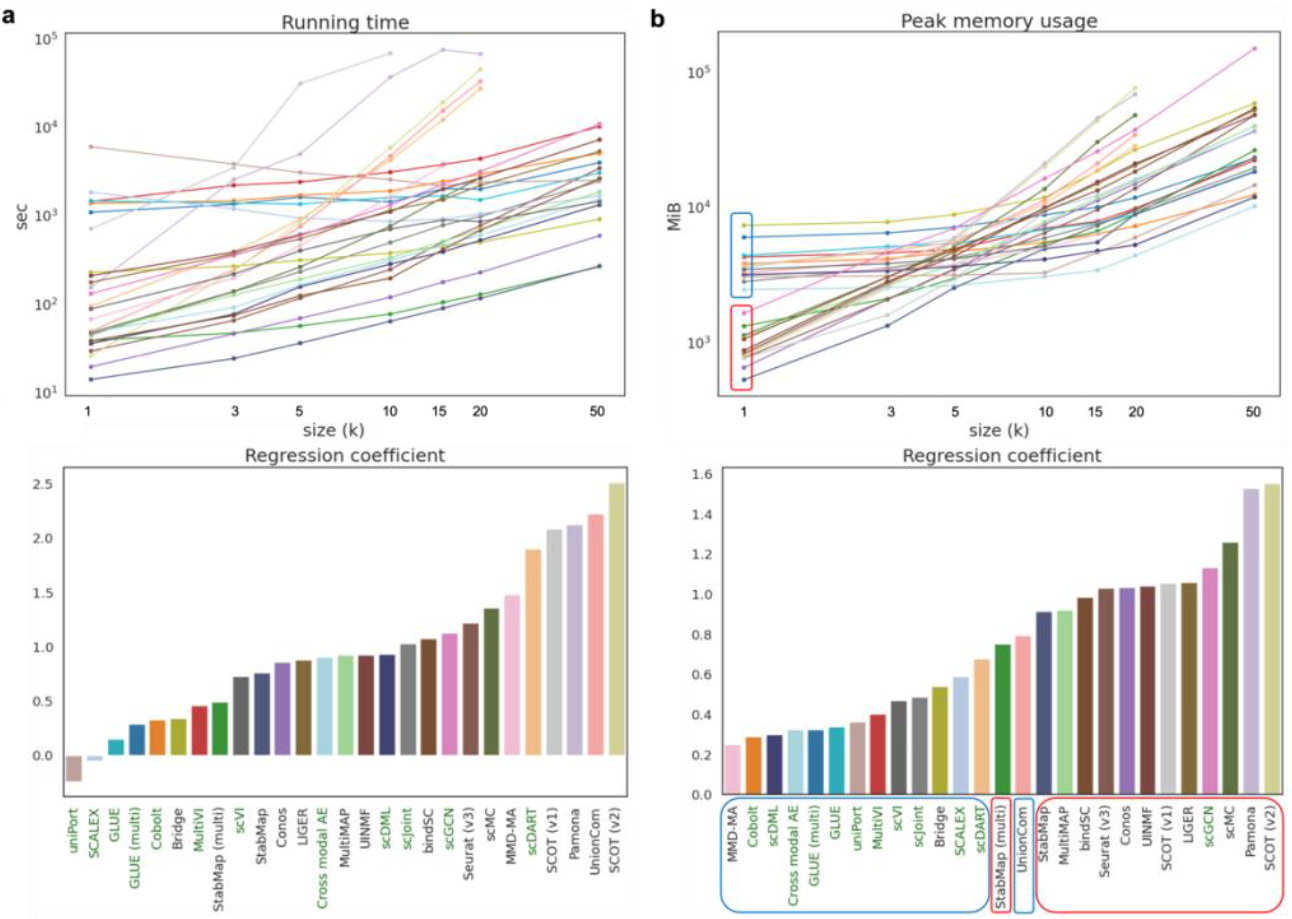
Computational efficiency comparison among 27 methods across 7 data sizes. (a) Running time comparison. (b) Peak memory usage comparison. The lineplot at the top of each subfigure shows the trends of the corresponding computational efficiency metric of all methods across increasing data scale. Both the y-axis and the x-axis are in log scale. The barplot at the bottom presents the coefficient of each method’s log-scaled metric fitted using linear regression on log-scaled data size. The color code for each method is consistently used in all plots. Methods whose names are in green are deep-learning-based methods. For (b), methods whose lineplots are marked by a red square at size = 1k are also highlighted by red squares along the x-axis of the barplot. So as for the methods whose lineplots are marked by a blue square. Heatmap view of running time and memory usage comparison across all methods and data sizes can be found in Supplementary Figure S14.

From Figure 5b, we can observe that the top five methods that had the fastest increase in peak CPU memory usage were SCOT (v2), Pamona, scMC, scGCN and LIGER. Among them, scGCN is a deep-learning-based method, but it involved a data processing step where both intra-data and inter-data graphs were constructed that consumed a lot of memory. The top five methods that scaled slowest as the data size increased were MMD-MA, Cobolt, scDML, cross-modal AE and GLUE (multi), among which the last four are all deep-learning-based methods and all of them used GPU for training. We found that if we divided all methods into two groups based on their peak memory usage at data size 1k (Figure 5b lineplot), all methods from the blue group except UnionCom were the top methods that scaled slowest with the data size (Figure 5b barplot). This suggests that the memory usage at a small data size was related to the increasing rate of a method’s memory usage when the data scaled up. Methods that consumed more memory at a relatively small sample size (1k) usually scaled better with larger data. Moreover, we found all methods except Bridge utilized GPU and Bridge relied on a technique called dictionary learning that is inherently scalable to large datasets [16].

## 3 Discussion

We performed a comprehensive benchmarking study on 27 computational tools that can be used to perform scATAC-seq label annotations, where five tools can directly perform label transfer from scRNA-seq to scATAC-seq data and 22 tools were originally designed for joint embedding (label transfer via training an additional kNN classifier), across 27 tasks composed of single-cell RNA and ATAC data collected from both human and mouse tissues. Moreover, we studied the impact of the data properties (data imbalance and cross-omics cell type dissimilarity) and method properties (utilization of paired data, peak information, and semi-supervised training strategy) on the performance of different methods. Prediction accuracy was evaluated on common cells and ATAC-specific cells separately. In addition, the running time and the peak CPU memory usage of each method were compared across a range of data sizes.

For the 18 tasks that did not involve additional paired data, the best performers in terms of prediction accuracy on common cells that were among the top three performers in more than half of the tasks were GLUE (16/18) and bindSC (13/18), followed by MultiMAP (6/18), UINMF (4/18), Seurat (v3) (4/18), scJoint (4/18) and uniPort (3/18). For the nine tasks that had available paired data from the same tissue, the methods that were among the top three performers in at least three tasks were Bridge, GLUE (multi), bindSC and GLUE. For ATAC-specific cells, scGCN was the one among all 27 methods that had the most similar performance to a random classifier, which was hypothetically the best classifier for ATAC-specific cells. Moreover, we found that scJoint had high enrichment scores and weighted accuracy, indicating its tendency to misclassify unobserved cell types with high confidence. The methods that consistently performed poorly were MMD-MA, Pamona, UnionCom, SCOT (both v1 and v2) and StabMap. Except for StabMap, all these methods relied on cell-cell distance information and tried to align such information from RNA and ATAC without incorporating any prior information about the relationships between features from the two modalities, causing potential misalignment of cell types of similar abundance instead of similar biology. For StabMap, although it related each ATAC peak with its nearby gene, it only kept one ATAC peak if multiple peaks were related with a gene and treated the ATAC peak and the gene as the same feature. This aggressive strategy might cause the loss of useful biological information.

Based on the observations from the 27 tasks, we concluded that when paired data from the same tissue were available to be jointly integrated with unpaired data, the best methods were Bridge and GLUE (multi) that could use such paired data. For mKidney, mBrain (atlas), mBrain (snap) and mBrain (10x), these are the tasks where neither Bridge nor GLUE (multi) was among the top two performers. The sequencing depths of their multimodal ATAC data were all relatively low with the median number of ATAC features per cell ranging from 570 to 1,086 (Supplementary Table S2). While for other tasks, the numbers were usually in thousands. Such high sparsity might be the reason why Bridge and GLUE (multi) performed relatively poorly on these tasks. When additional paired data were not available, the best performers were bindSC and GLUE and a common property of these two methods was that they both used the ATAC peak data instead of purely relying on gene activities. To align the feature sets between RNA gene-level data and ATAC peak-level data, GLUE utilized a prior knowledge graph that recorded the regulatory interactions among genes and ATAC peaks, and bindSC used a technique call bi-order canonical correlation analysis to align both the cells and features between RNA and ATAC data with gene activity matrix only used for algorithm initialization.

Even though GLUE (multi) and Bridge could utilize paired data and achieved the best performance on the third group of tasks, we found the utilization of paired data did not always improve the prediction accuracy, such as MultiVI versus scVI and StabMap (multi) versus StabMap. This might be due to some inherent limitations in the design of these algorithms. For example, MultiVI calculated the joint embedding from two modalities by just averaging the modality-specific embeddings and StabMap only used one dataset as reference and performed linear transformation to map other datasets to the reference.

Moreover, we found the introduction of semi-supervised strategy had a negative impact or no effect on most methods. The method that was most negatively impacted by this was GLUE. Semi-supervised training did increase the prediction accuracy of scJoint on average, but the change was not significant. The only significant and positive change was observed on scDML, which is a method based on deep metric learning by first finding similar and dissimilar clusters between RNA and ATAC data. Note that the semi-supervised training was realized through concatenating an additional layer of neural net as a classifier to the embedding layer of a tested method. Therefore, the observation that semi-supervision only worked on scDML might be because both are clustering-based techniques. This suggests that semi-supervised training is not always compatible with a method.

Apart from the choice of the computational tool, we found that the prediction accuracy can be affected by the properties of a dataset. Basically, datasets with large proportions of modality-specific cell types, high discrepancy in the common cell type compositions between modalities, and low cross-omics similarity in common cell types had poor accuracy scores across all the methods.

Almost all the methods were negatively impacted by the data imbalance even though some methods tried to deal with discrepancy in cell type distributions between two modalities through their unique designs, such as using mutual nearest neighbors [12-16], reweighting cells [31], unbalanced optimal transport [22, 27, 28] or integrative non-negative matrix factorizations [17, 33]. Since we removed imbalance at two levels by only removing modality-specific cell types and by making cell types perfectly balanced between RNA and ATAC, we were able to perform pairwise comparisons among these two experiments and the original results. We found some methods (e.g. scJoint, bindSC, UINMF, LIGER and Conos) could only benefit from removing modality-specific cell types, which suggests their algorithms can deal with the imbalance in cell type compositions as long as cell types from two modalities are the same. Some methods (e.g. Bridge, GLUE, MultiVI, Seurat (v3), MultiMAP, scGCN and SCALEX) could benefit further from balancing the cell type compositions between modalities. This implies that these algorithms cannot properly deal with data imbalance.

Although data binarization was adopted by scJoint as a potential way to reduce the difference in data distribution between two modalities, we found the prediction accuracy was negatively affected and the F1 of entropy and enrichment was positively affected by data binarization for most methods. We speculate that this was because binarization removed the differences among cell types within each modality and thus made it harder for algorithms to label cell types. Another potential reason was that some methods assumed that single-cell data followed some over-dispersed distributions like negative binomial distribution, so binarization could have caused the violation of this model assumption.

In terms of method scalability, we found the most time and memory-consuming methods are those that involved matrix optimization (e.g. SCOT, Pamona and UnionCom) or pairwise distance calculation among all cells (e.g. scMC, scGCN and scDART). In contrary, the most memory-efficient methods were usually the deep-learning-based ones (e.g. uniPort, SCALEX, GLUE, Cobolt and MultiVI) that could use GPU and minibatch training strategy. The methods that consumed less time were also deep-learning-based algorithms, especially those that adopted early stopping and fixed number of training iterations. Iteration-based training can largely save the running time compared to epoch-based training given a fixed minibatch size. This is because for an epoch all cells in the training data are used, while for an iteration only a minibatch of cells are used. Apart from deep-learning-based method, we found Bridge was also both time and memory efficient because it utilized dictionary learning and only performed heavy computation on a subset of data.

Our study has some limitations. First, although we tried to collect as many data as possible for this benchmarking study, the types of tissues and sequencing technologies we covered were still limited, especially for tasks that had both unpaired data and corresponding paired data. Second, we used the cell type labels provided by their original publications as the ground truth. However, the task of cell type annotation is usually a combination of results from computational tools and manual adjustment by experts, so the derived labels could be biased and subjective. For some finer levels of cell type annotation, even researchers from the same field can have different opinions.

To summarize, we have the following suggestions for users to choose among currently available tools. If high quality paired data can be found to transfer labels from unpaired scRNA-seq to scATAC-seq data, Bridge and GLUE (multi) are likely the best method; otherwise, bindSC and GLUE are the recommended methods. Both Bridge and bindSC are methods written in R and compatible with the most popular single-cell R package Seurat, so R users should feel more comfortable to use these two methods. For Python users, GLUE can be a better choice not only because it is written in Python and compatible with Python’s single-cell module Scanpy [66], but also because this tool is able to run both with and without pairing information between two modalities. Note that none of these methods had a good performance on ATAC-specific cells. Therefore, if one cares about ATAC-specific cell types, a better strategy might be using scGCN first to identify potential cells unique to ATAC, and manual annotations can be performed on cells that have high entropy and low enrichment.

For future directions, we think an ideal method for label transfer from scRNA-seq to scATAC-seq should take the following aspects into consideration. First, such a method should thoroughly deal with the data imbalance between scRNA-seq and scATAC-seq data. Second, such a method should take care of the peak-level information instead of purely relying on gene activity matrix to represent the ATAC data. Third, it should be able to properly incorporate paired data if such data with high quality are available. Fourth, it should not over-normalize the original data, like through data binarization. Some studies have argued that the binarization of scATAC-seq data would suffer from loss of information [67, 68]. Fifth, semi-supervised training strategy should be used with caution as this would not always benefit a method and even could have a negative effect. Last, future methods can borrow the idea from GLUE that includes prior knowledge to inform the relationships among features from different modalities.

## 4 Materials and Methods

### 4.1 Single-cell data preprocessing

A full list of data used in this study can be found in the Supplementary Table S1. For each tissue, it can contain at most three types of datasets, namely unimodal RNA data, unimodal ATAC data and multimodal data (RNA and ATAC measured simultaneously for the same cell). To facilitate the evaluation of label prediction performance, we manually unified the naming conventions of cell labels provided in the scRNA-seq and scATAC-seq. We also removed cell types from each dataset that contained less than 10 cells. As a common practice, for all the scATAC-seq data (both unimodal and multimodal), if the raw gene activity matrix in integers was not available, we calculated it using function ‘GeneActivity’ in R package Signac with the fragment files as inputs [69]. For all the tissues that had all types of data, we were able to use them to benchmark methods that could integrate paired and unpaired data together, including Bridge, MultiVI, Cobolt, GLUE (multi) and StabMap (multi). Therefore, we had to align the peak set of unimodal ATAC and that of the multimodal ATAC data. If the fragment files of the multimodal ATAC data were available, we requantified the abundance of multimodal ATAC peaks using the unimodal peak set as reference by the ‘FeatureMatrix’ function in Signac. As the last step of data processing, if a dataset contained more than 20k cells, we downsampled it proportionally to contain 20k cells except for the cell types that originally contained less than 100 cells. Descriptions of preprocessing pipelines specific to each dataset, like exceptions to the common practices mentioned above, are provided below. Details for data preprocessing can be found in our GitHub repository. Tissues that only have unimodal data (unpaired) or multimodal data (paired) are indicated correspondingly in the following paragraphs.

#### Human T-cell-depleted bone marrow (TDBM) (paired)

There was no discrepancy between the analysis pipeline we performed for this tissue and the one described previously.

#### Human kidney (unpaired)

There was no discrepancy between the analysis pipeline we performed for this tissue and the one described previously.

#### Human fetal atlas (unpaired)

There was no discrepancy between the analysis pipeline we performed for this tissue and the one described previously. For this tissue, the raw gene activity matrix was available online.

#### Human basal cell carcinoma (BCC) (unpaired)

There was no discrepancy between the analysis pipeline we performed for this tissue and the one described previously.

#### Human heart (unpaired)

There was no discrepancy between the analysis pipeline we performed for this tissue and the one described previously.

#### Human PBMC COVID (unpaired)

For the scATAC-seq data, although the original publication claimed that the reference genome they used was hg19, it was actually hg38 because only hg38 could yield correct TSS enrichment score. Therefore, we used hg38 to calculate gene activities for ATAC.

#### Human PBMC

The reference genomes used for the unimodal ATAC (hg19) and the multimodal ATAC (hg38) data were different and only the latter had public raw sequence data in fastq formats. We remapped the multimodal data using cellranger-arc to hg19 to get the peak count matrix and fragment files to align peak sets between the unimodal and the multimodal ATAC data.

#### Human BMMC

This is so far the largest single-cell multimodal RNA and ATAC dataset with well-annotated labels and hierarchical batch structures. To mimic the case where scRNA-seq, scATAC-seq and multimodal data were measured separately, we manually separated all batches to three groups without any overlaps. Specifically, batches s1d2, s1d3, s3d3, s4d9, and s3d10 were used as scRNA-seq (26,450 cells), s2d4, s2d5, s3d6, s3d7, and s4d8 were used as scATAC-seq (24,332 cells), and s1d1, s2d1, s4d1 were used as multimodal data (18,467 cells).

#### Human HSPC

For the unimodal ATAC data, we didn’t use Signac to calculate the gene activity matrix because we couldn’t find any accompanying fragment files which were required by Signac to do the calculation. Therefore, we wrote a function called ‘CreateGeneActivityMatrix’ (can be found in our online GitHub repository) that can calculate gene activities purely rely on the peak count matrix by summing up peak counts within the 2000 base pairs around the transcription starting site of a gene. For the multimodal ATAC data, the gene activity calculation was still performed by default using Signac because fragment files were available. Moreover, since the original reference genomes used for the unimodal ATAC (hg19) and the multimodal ATAC (hg38) data were different and no raw fastq files were found, we lifted peaks of the unimodal ATAC data over from hg19 to hg38 using UCSC liftover utility for peak set alignment.

#### Mouse skin (paired)

There was no discrepancy between the analysis pipeline we performed for this tissue and the one described previously.

#### Mouse retina (paired)

For the raw gene activity matrix, we used the one provided by the original publication, which was calculated by Cicero.

#### Mouse spleen (unpaired)

For the raw gene activity matrix, we used the one provided by the original publication, which was calculated by snapATAC.

#### Mouse endothelial (unpaired)

There was no discrepancy between the analysis pipeline we performed for this tissue and the one described previously.

#### Mouse atlas (unpaired)

There were two types of RNA data: one was sequenced by scRNA-seq (droplet) and the other was sequenced by SMART-seq (FACS). We processed and benchmarked these two types of RNA data separately with the same ATAC data. For the ATAC data, we used the gene activity matrix provided by the original publication.

#### Mouse brain

We had three pairs of unimodal RNA and ATAC data for mouse brain. The first pair was composed of the brain cells from the previous mouse atlas data. For the RNA data, only the FACS data were used because the droplet data didn’t have any brain cells. For the rest two pairs, they shared the same RNA data sequenced by Drop-seq. The two unimodal ATAC data were from separate studies, one sequenced by scATAC-seq and the other sequenced by snATAC-seq. For the former one, we used the default method (Signac) to calculate the gene activity matrix; for the latter one, we used the provided raw gene activity matrix calculated by snapATAC. All three pairs of unimodal data shared the same multimodal data from SNARE-seq. Since the reference genome used for the mouse atlas ATAC data (mm9) was different from the one used for the multimodal ATAC data (mm10), we lifted peaks of the mouse atlas brain ATAC data over from mm9 to mm10 using UCSC liftover utility for peak set alignment.

#### Mouse primary motor cortex (MOp)

The raw gene activity matrix for the unimodal ATAC data was extracted from the snap objects provided by the original publication, which was computed using snapATAC.

#### Mouse embryo

For the unimodal ATAC data, although the fragment files were not available, we downloaded the fastq files and used the pipelines provided by the authors on GitHub (https://github.com/BPijuanSala/MouseOrganogenesis_snATACseq_2020) to generate a BAM file. After indexing the BAM file using SAMtools, we used sinto to generate a fragment file and then used tabix to sort, block-gzip compress (bgzip) and index the file.

#### Mouse kidney

For this tissue, we requantified the unimodal ATAC peaks on the multimodal peak set, because only the fragment files of the unimodal ATAC data were available. Moreover, like how we calculated the gene activities for human HSPC’s unimodal ATAC data, we calculated gene activities for the multimodal ATAC data using the raw peak count data via ‘CreateGeneActivityMatrix’ because no fragment files were available.

### 4.2 Description and implementation of methods

#### Conos

Conos is designed as a graph-based batch effect removal method. The joint graph embedding using nearest neighbors and Pearson correlation is constructed as the first step to connect all cells. Then, the label transfer from reference data to query data can be implemented by information propagation between graph vertices through an iterative diffusion process.

#### Seurat (v3)

Seurat first identifies a set of anchors between the reference and the query data through canonical correlation analysis (CCA) and mutual nearest neighbors (MNNs). Then, a weight matrix is constructed to quantify the distance between each query cell and anchor cell in the query data by a Gaussian kernel. Last, the prediction score of any cell in the query data is calculated as a weighted average of labels of anchor cells in the reference data. Both the gene activity and raw peak count matrices are needed, with the former matrix required to obtain cross-omics anchors through CCA and the latter one recommended for nearest neighbor construction among the ATAC cells.

#### scGCN

The first step of scGCN is to build a hybrid graph of all cells using MNNs approach and CCA. Based on the constructed graph, a semi-supervised graph convolutional neural network is trained to embed cells from both reference and query data on the same latent space and predict cell type labels for cells in the query data.

#### scJoint

Like scGCN, is a semi-supervised neural network trained to jointly embed cells from both scRNA-seq and scATAC-seq. Different from scGCN that directly utilizes the trained network to predict probability vectors through Softmax layers, scJoint performs label transfer by training an additional kNN classifier in the embedding space. The loss function of scJoint is composed of a dimensionality reduction loss, a cosine similarity loss and a cross-entropy loss (calculated only using scRNA-seq data and the known cell labels). The role of the cosine similarity loss is to maximize the similarity between best aligned RNA and ATAC cells in the latent space. To account for the case where RNA and ATAC do not share the same cell types, only the top 80% of cells with the highest cosine scores are used to calculate this loss term.

#### Bridge

This method utilizes multimodal data as a bridge to transfer labels from scRNA-seq to scATAC-seq. The multimodal dataset is treated as a dictionary and each cell is an atom, on which dictionary representations of both unimodal scRNA-seq and scATAC-seq are constructed. After dimensionality reduction of multimodal cells via Laplacian Eigendecompostions, unimodal cells can be embedded on the same space by the dictionary representations. Then, the final label transfer can be achieved by any single-cell integration techniques and Bridge chooses mnnCorrect.

### All methods below are joint embedding methods for single-cell multi-omics data that do not directly perform label transfer from RNA to ATAC

#### MMD-MA

This is a manifold alignment method for unpaired single-cell multi-omics datasets based on maximum mean discrepancy (MMD). It aligns and embeds two datasets in Reproducing Kernel Hilbert spaces by minimizing the MMD across datasets. It does not require any overlap between the feature space of two datasets.

#### UnionCom

This method is based on metric space matching to jointly embed unpaired single-cell multi-omics datasets to a latent space. Specifically, it aligns cells across omics by matching the distance matrices by matrix optimization. Like MMD-MA, it does not require any correspondence information among features of the two datasets, so gene activity calculation is not needed.

#### SCOT

SCOT is an unsupervised alignment tool based on optimal transport (OT) for unpaired single-cell multi-omics data integration and it does not require any correspondence information among either cells or features. As the first step, an intra-domain distance matrix is calculated for each data by constructing a kNN graph. Then, cell-to-cell correspondence probabilities are derived based on Gromov-Wasserstein OT to align datasets cross omics. There are two versions of SCOT, for version 1, it cannot handle the situation where two omics have disproportionate cell type representations as it is based on global OT; For version 2, SCOT is extended to handle such case using unbalanced OT. Another difference is that version 1 integrates data without changing the number of input dimensions, but version 2 embeds data by projecting them to a joint latent space.

#### Pamona

Like SCOT, this method also relies on OT and is a partial Gromov-Wasserstein-based manifold alignment algorithm that can handles differences in cell type compositions between two modalities by allowing only a fraction of the total mass to be transported.

#### MultiMAP

This algorithm is designed based on the idea of UMAP to integrate and embed single-cell multi-omics datasets to a two-dimensional space. It first calculates both the intra and inter omics geodesic distances among cells and then constructs a multi-omics neighborhood graph based on these distances. Finally, it projects the data into a low-dimensional space by minimizing the cross-entropy of the graphs in the latent space and the manifold space. Both original peak data and gene activity data from ATAC are used to calculate intra and inter omics cell distances respectively.

#### scVI

This is a deep learning method based on a variational autoencoder (VAE) to integrate scRNA-seq data from different batches. Batch information is provided as a categorical variable to both the encoder and decoder to help derive a batch-free latent embeddings of cells from all datasets. Here, we applied scVI to jointly embed unpaired scRNA-seq data and scATAC-seq data’s calculated gene activity matrix.

#### Cross-modal AE

This is an autoencoder-based method for unpaired multimodal data integration and embedding. After projecting data into a latent space using modality-specific encoders, data are aligned through adversarial training by connecting a discriminator to the latent space.

#### SCALEX

Like cross-modal AE, SCALEX is a deep learning method based on a VAE for single-cell data integration and can be applied to jointly embed single-cell RNA and ATAC data if gene activity matrix is calculated from the scATAC-seq data. It projects data into a batch-invariant latent space by using a batch-free encoder and a batch-specific decoder.

#### scDML

This method is also a deep learning method but utilizes deep metric learning to remove batch effects among datasets in a latent space projected through an encoder. Initial clusters are inferred by algorithms like Louvain before training a neural network and both intra and inter data cluster similarities are assessed by computing the number of kNN and MNN pairs, respectively. Deep metric learning is used to ensure that similar clusters among datasets are merged and unsimilar clusters are kept away from each other in the latent space.

#### uniPort

This method can perform joint embedding of unpaired single-cell multi-omics datasets by combining a coupled VAE and minibatch unbalanced OT. It utilizes the information from both common highly variable genes and data-specific genes by incorporating multiple decoders.

#### scDART

This is a scalable deep learning method that is designed specifically for integrating unpaired scRNA-seq and scATAC-seq data in a low-dimensional space. Unlike most methods, scDART doesn’t require a pre-calculated gene activity matrix for the ATAC data. Instead, it introduces a non-linear gene activity function module that connects ATAC peaks (first layer) with RNA genes (second layer) and the weight matrix of this module is learned by putting a binary mask on it. The mask matrix is derived based on the genomic locations of peaks and genes. Additionally, scDART utilizes MMD as a kernel-based discrepancy to align similar cells across modalities in the latent space.

#### GLUE

Like some previous deep learning-based methods, GLUE is based on VAE, but the difference is that in addition to using omics-specific VAEs to learn cell embeddings, GLUE incorporates a knowledge-based graph VAE that utilizes prior information about the regulatory interactions among genes and ATAC peaks. Therefore, GLUE is immune to the information loss by avoiding the calculation of gene activities and the cross-omics joint embeddings can be guided by the prior knowledge graph. Moreover, a discriminator is used to align cell embeddings from different omics through adversarial training. To deal with the case where cell type compositions differ among omics, GLUE introduces weighted adversarial alignment that assigns weights to cells to balance cell distributions across omics. GLUE also has a extend version that can be used when both paired and unpaired data are available by penalizing distances between paired cells in the latent space.

#### scMC

This method relies on variance analysis to deconvolutes technical and biological variations among different single-cell datasets at the cell cluster levels. It starts with the identification of putative cell clusters for each data and then the inference of shared clusters between two datasets. Variance analysis is used to derive the correction vectors for batch effect removal.

#### bindSC

This method is based on bi-order canonical correlation analysis (CCA) for unpaired single-cell multi-omics data integration. CCA is performed iteratively to align both cells and features from two modalities. bindSC requires both the original peak matrix and calculated gene activity matrix from scATAC-seq data, but the gene activity matrix is only used to initialize the feature matching.

#### LIGER

This method relies on integrative non-negative matrix factorization (iNMF) to integrate and embed unpaired single-cell multi-omics datasets by delineating shared and data-specific features (metagenes). This method requires sharing features across datasets, so gene activity calculations are needed when integrating scATAC-seq and scRNA-seq data.

#### UINMF

UINMF is an extension of LIGER for the case where features are partially shared across datasets (but there must exist a set of features shared across all datasets). The key idea is to let data-specific features to inform the factorization by introducing an additional metagene matrix. To apply UINMF, both the original peak data and gene activity data from ATAC should be provided along with the RNA gene data so that both unshared and shared features from ATAC can be used to inform the joint embedding.

#### MultiVI

MultiVI is built based on scVI that can integrates both paired and unpaired scRNA-seq and scATAC-seq data so that gene activity calculations are not needed. The distance between two modalities is penalized through minimizing the KL divergence between their distributions in the latent space and the joint embeddings are calculated as the average of the embeddings from two modalities.

#### Cobolt

Like MultiVI, Cobolt is a multimodal VAE (MVAE) framework that can be used to jointly embed unpaired single-cell RNA and ATAC data by using additional paired multi-omics data. To better align data from different modalities in the latent space, Cobolt first fits the MVAE model and then uses the paired data to train cross-omics predictors that can correct for missing modalities of the unpaired data.

#### StabMap

StabMap is designed for mosaic data integration like UINMF where features are partially overlapped. However, it is different from UINMF and is more generalized in the sense that no set of features shared across all datasets is required. The only requirement is that there is a way to draw a path from every data to every other data and two data are connected if there exist shared features. Therefore, StabMap can be used to integrate unpaired scATAC-seq and scRNA-seq data with paired data without calculating gene activities. According to the original paper, StabMap can also be applied to unpaired data only, by linking ATAC peaks with nearby genes and treat them as the same features.

For all the joint embedding methods that do not explicitly perform label transfer, we first obtained the latent representations of both scRNA-seq and scATAC-seq data using the default or recommend number of latent dimensions for each method. When no default or recommended number is available, we used 20. Then, a kNN classifier (k=30) was trained using the latent representations and known cell labels of scRNA-seq data. Last, we used the trained kNN classifier to get the predicted probability matrix for the corresponding scATAC-seq data for all downstream evaluations.

Note that for StabMap (base version without paired data being used), scDART and GLUE, they all require the region-to-gene relationships to link ATAC peaks and RNA genes as prior information. We used GLUE’s Python function ‘scglue.genomics.rna_anchored_guidance_graph’ to obtain the required format of region-to-gene relationships for all the three methods. Since the reference genome is needed when running this function, we used the genome version indicated in the original publications of each scATAC-seq dataset (shown in Supplementary Table S1).

For methods that do not require additional paired multi-omics data, we used the raw count matrix of scRNA-seq and raw count matrix or gene activity matrix or both of scATAC-seq data based on the requirements of each method (see Table 1) as inputs. For methods that require paired data to integrate or transfer labels from unpaired RNA data to ATAC data, we provided raw count matrices of scRNA-seq, scATAC-seq and multimodal data (with the peak sets of unimodal and multimodal ATAC data aligned). The implementation of each method followed the instructions on their websites or in their original publications. Details can be found in the scripts on our GitHub repository and package versions can be found in Supplementary Table S3.

### 4.3 Evaluation metrics

We assessed the performance of different cross-omics label transfer methods from two aspects, namely accuracy and scalability. For prediction accuracy evaluation, we divided ATAC cells into two parts and designed metrics for them separately: (1) cells whose cell types also existed in the corresponding scRNA-seq data and we call these common cells; (2) cells whose cell types were only observed in scATAC-seq data and we call these ATAC-specific cells. For scalability evaluation, we compared both the running time and memory usage among all methods across selected gradients of sample sizes (see details in 4.4 Benchmarking design).

#### Overall accuracy

After getting the predicted probability matrix across all cells in scATAC-seq, the cell type that had the highest predicted probability was assigned to each cell as the predicted label. Then the overall accuracy was calculated using the predicted labels and true labels for all common cells. We didn’t calculate this metric for ATAC-specific cells because it would be exactly zero.

#### Balanced accuracy (macro recall)

The overall accuracy can be easily affected by the composition of cell types in data. To remove that influence and treat all cell types equally, we calculated balanced accuracy for all common cells, which is the average of recalls for each cell type (macro recall),

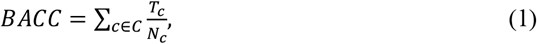

where *C* is the set of all common cell types, *N*_*c*_ is the number of cells in cell type *c*, and *T*_*c*_ is the number of cells correctly predicted as cell type *c*.

#### F1 score

Precision is defined as true positive (TP) over the summation of TP and false positive (FP) and recall is defined as TP over the summation of TP and false negative (FN). F1 score is the harmonic mean of precision and recall,

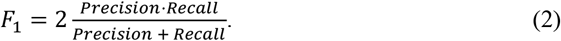

Since this is a multi-class classification problem, we need to specify whether we want macro or micro level metrics. It is easy to show that overall accuracy is equivalent to micro precision, recall and F1 score under the multi-class scenario. Therefore, we calculated macro level precision and recall in this study, which is the average of precisions and recalls obtained for each class. Then, macro F1 score is calculated based on macro precision and recall. Like overall accuracy and balanced accuracy, F1 score was calculated only for common cells.

#### Weighted accuracy

To account for the prediction uncertainty and similarity across cell types. We proposed a weighted accuracy (WACC) by taking the average of the predicted probability vector weighted by cell type similarities:

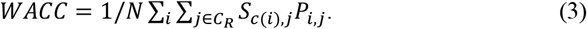

In the equation above, *P* is the predicted probability matrix with each row as a cell in scATAC-seq and each column as a cell type observed in scRNA-seq reference data. *C*_*R*_ is the set of all cell types in scRNA-seq and *N* is the total number of scATAC-seq cells. *S* is a cross-modality cell type similarity matrix with each row as a cell type in scATAC-seq and each column as a cell type in scRNA-seq and *c*(*i*) is a function mapping cell *i* to its true cell type label. Since for ATAC-specific cell types, their similarities to cell types in RNA data could be assessed through common cell types and recorded in the cross-modality similarity matrix *S*, weighted accuracy was calculated on both common cells and ATAC-specific cells.

The similarity matrix was calculated in three steps. First, partition-based graph abstraction (PAGA) [70] was performed on the normalized count matrix of scRNA-seq and gene activity matrix of scATAC-seq separately. Then, the within-modality similarity matrix was calculated based on the Euclidean distance of each pair of cell types using the PAGA positions. For cell types *i* and *j*, their within-modality similarity was calculated as:

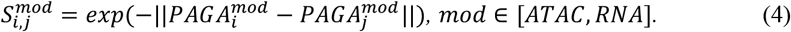

Last, we calculated the cross-modality similarity matrix using the two within-modality matrices by considering three scenarios. If two cell types existed in both modalities, their similarity was calculated as the average of two within-modality similarities:

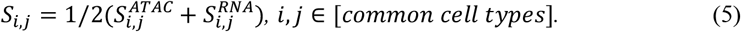

If one cell type is modality-specific, its similarity with any common cell type would be the similarity calculated using the modality that contained the two cell types:

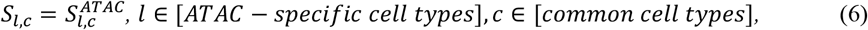

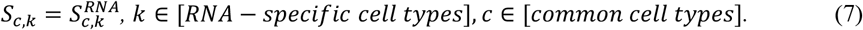

If a cell type *l* only existed in scATAC-seq and the other cell type *k* was only observed in scRNA-seq, their similarity was calculated as

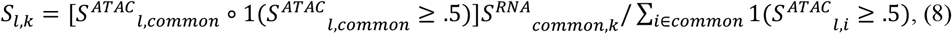

where *S*^*ATAC*^ and *S*^*RNA*^ are within-modality similarity matrix for ATAC and RNA, respectively, 1 represents an indicator function and *common* is the set of all common cell types. The first product is Hadamard product which is element wise and the second product is matrix multiplication. This metric was calculated for both common cells and ATAC-specific cells separately.

#### Entropy and enrichment

To further evaluate the performance of methods on ATAC-specific cell types, we borrowed the two metrics proposed in scGCN which are scaled entropy and enrichment [14]. Scaled entropy is defined as

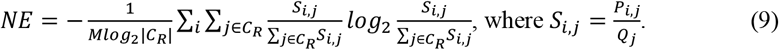

*P*_*i,j*_ is the predicted probability for cell *i* with unique cell type label in scATAC-seq and cell type *j*, and *Q*_*j*_ is the proportion of cell type *j* in scRNA-seq as the background probability. *C*_*R*_ is the set of all cell types in scRNA-seq and *M* is the total number of scATAC-seq cells with unique cell labels. The final score is normalized by *log*_2_|*C*_*R*_| to make it in the range of [0, 1]. Another metric is enrichment score,

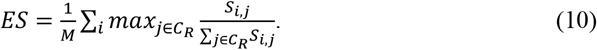

The enrichment score is also bounded within 0 and 1. For cell types only observed in scATAC-seq, an ideal method should deliver high normalized entropy and low enrichment score. Therefore, we also calculated an F1 score to combine these two

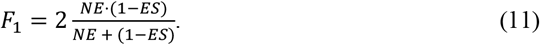

#### Running time and memory

All methods were run on Yale’s high performance computing clusters with one computing core. For methods that require GPUs as indicated in Table 1, they were run using GPUs; and for the rest methods, they were run using CPUs. The CPU of our device is Intel ® Xeon ® Gold 6240, 2.6 GHz, and the GPU is NVIDIA A100 with 40 GB RAM. When evaluating running time, we did not count the time used for data preprocessing (e.g. remaping to alternative reference genome, requantifing scATAC-seq peaks, and calculating gene activity matrix) because the needed steps for different tissues were different. For memory assessment, we used the peak memory usage to compare across different methods. To obtain the peak memory usage, we used ‘memory_profiler.memory_usage’ for Python methods and peakRAM for R methods.

### 4.4 Benchmarking design

In the following paragraphs, we will describe in detail the benchmarking design for experiments conducted in this study.

#### General performance comparison across different tissues

We ran all 27 methods, including 22 methods for unpaired data and 5 methods that involved additional paired data, on 27 dataset combinations, including 18 unpaired data (9 with available paired data from the same tissue) and 9 paired data. For the 9 paired datasets, they were used as if the RNA and the ATAC data were sequenced separately. Therefore, we ran in total 639 method and data combinations (22*27+5*9) to assess the general performance of all selected methods across different tissues.

#### Impact of cell type imbalance between modalities

The original RNA and ATAC data could contain different cell types and the compositions of cell types could vary significantly as well. We studied the impact of cell type imbalance between the two modalities on model performance, we manually subsampled the RNA and ATAC data for each tissue in two ways. First, we deleted all modality-specific cell types for each tissue to make the RNA and ATAC data contain the same set of cell types. We called this common cell type experiment. Second, apart from deleting modality specific cell types, we downsampled cells to make sure that the numbers of cells that belong to the same cell types are the same between RNA and ATAC. We called this balanced cell type experiment.

#### Impact of data binarization

We investigated whether data binarization could affect the performance of any methods across datasets we collected by binarizing both the RNA data and the ATAC data first before inputting them into the algorithms.

#### Effect of semi-supervised strategy

To investigate the effect of introducing semi-supervised training strategy to deep-learning-based methods on the performance of cross-omics label transfer. We downloaded and modified the original code of selected methods, including scDART, scDML, SALEX, cross-modal AE, GLUE, uniPort, scVI, MultiVI and Cobolt. Specifically, we connected the latent layer of each method with an additional linear layer whose number of nodes was equal to the number of clusters in the RNA data and added a cross-entropy loss calculated using the RNA data only to the original loss function. For scJoint, since it already employed the semi-supervised training strategy, we removed the cross-entropy term from its loss function to obtain an unsupervised version of scJoint.

#### Time and memory usage comparison

To study the scalability of all 27 methods. We used the mouse MOp data to measure the running time and peak memory usage across seven data sizes, including 1k, 3k, 5k, 10k, 15k, 20k, and 50k cells.

### 4.5 Data availability statement

All the single-cell data used in this study are publicly available. Detailed information of each data and their downloadable links can be found in Supplementary Table S1. The related scripts for reproducing results in this study will soon be available at our GitHub repository.

