## Supplementary Figures for "A Comprehensive Benchmarking Study on Computational Tools for Cross-omics Label Transfer from Single-cell RNA to ATAC Data"

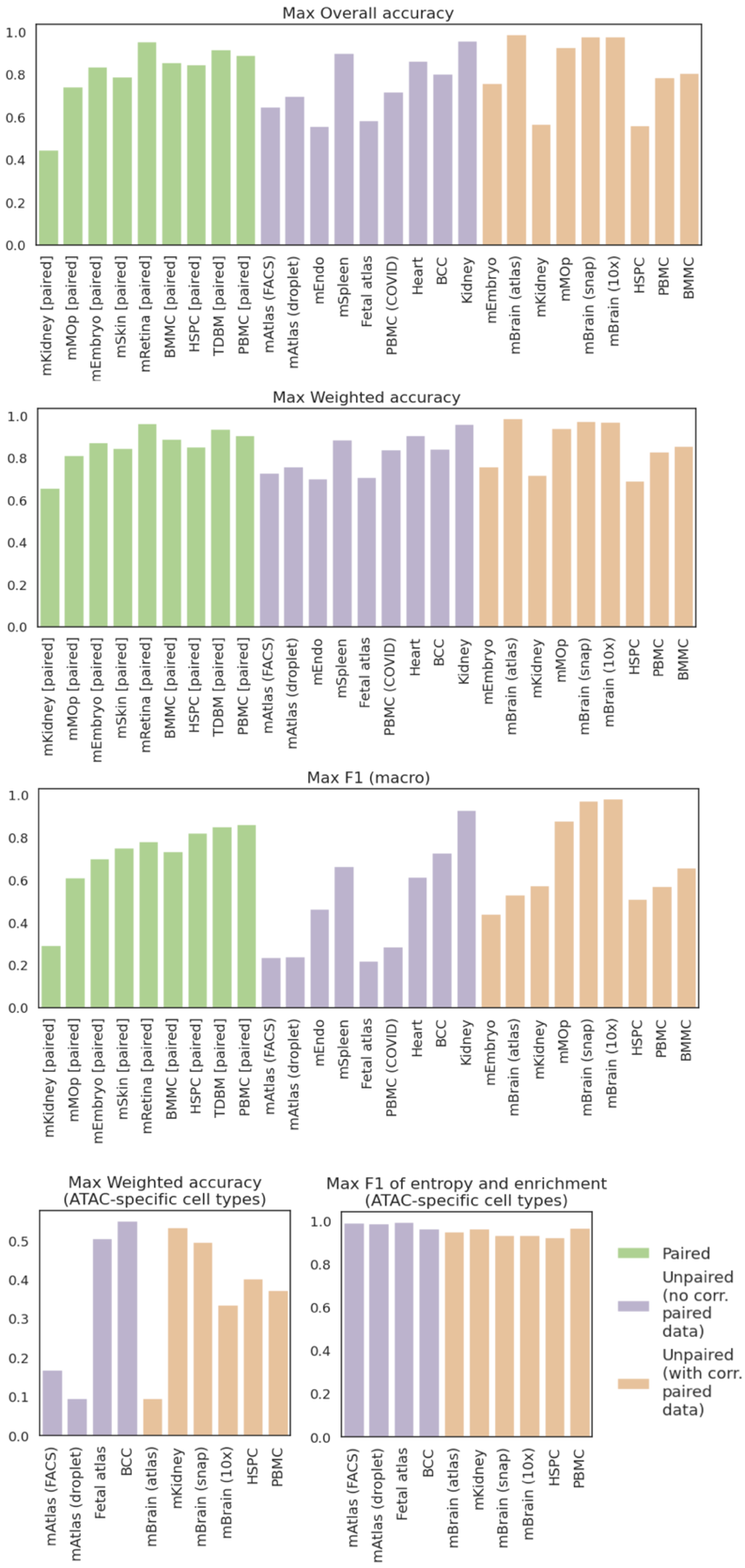

**Figure S1.** Supplementary information of the maximum values achieved on each task (dataset) across all the methods of metrics not shown in Fig. 3a. The order of datasets is the same as that in Fig. 3a for the same group.

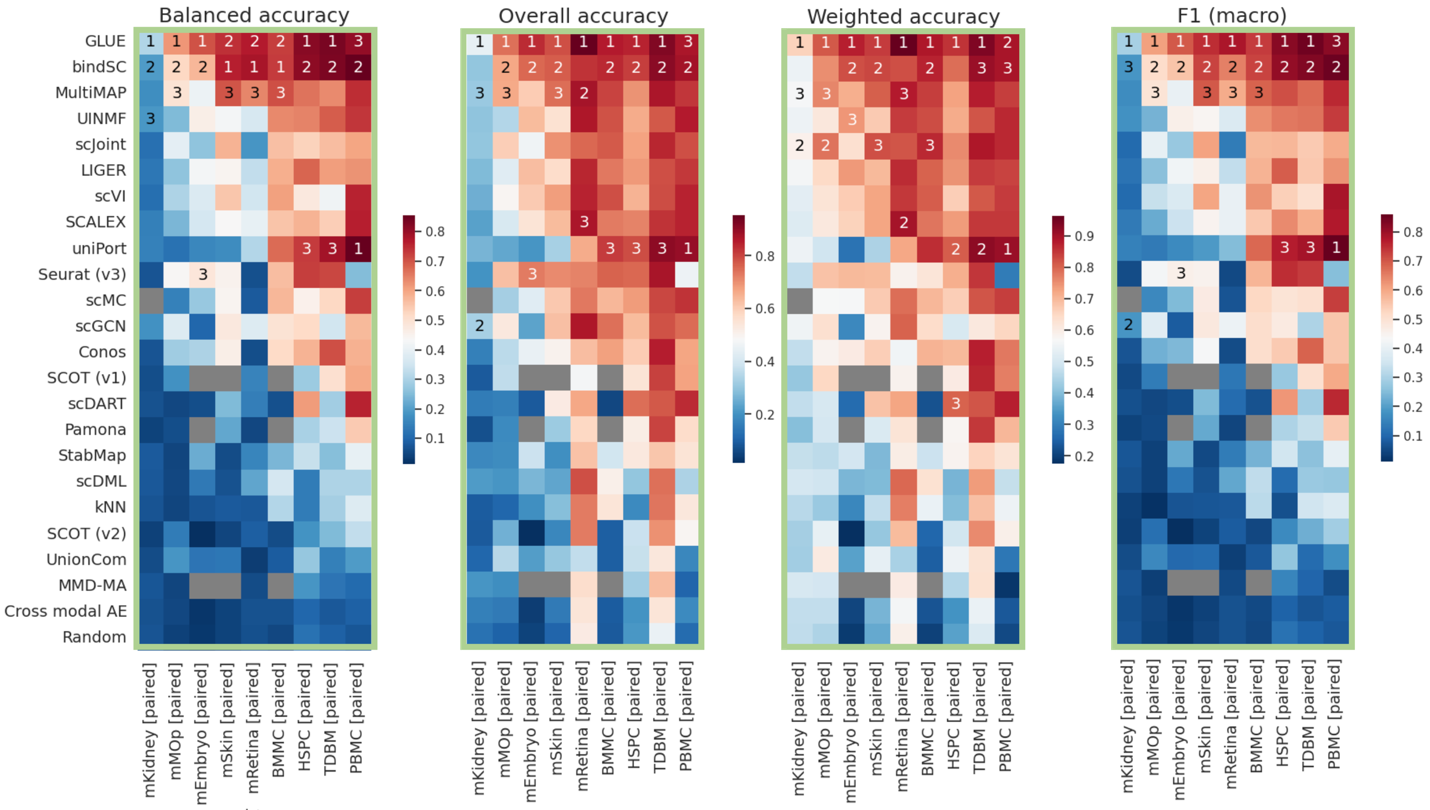

**Figure S2.** Detailed metrics for paired datasets (pairing information not used). For each heatmap, methods (rows) are placed in the same order as that in Fig. 2a and methods ranked top three in each task are marked by Arabic numbers.

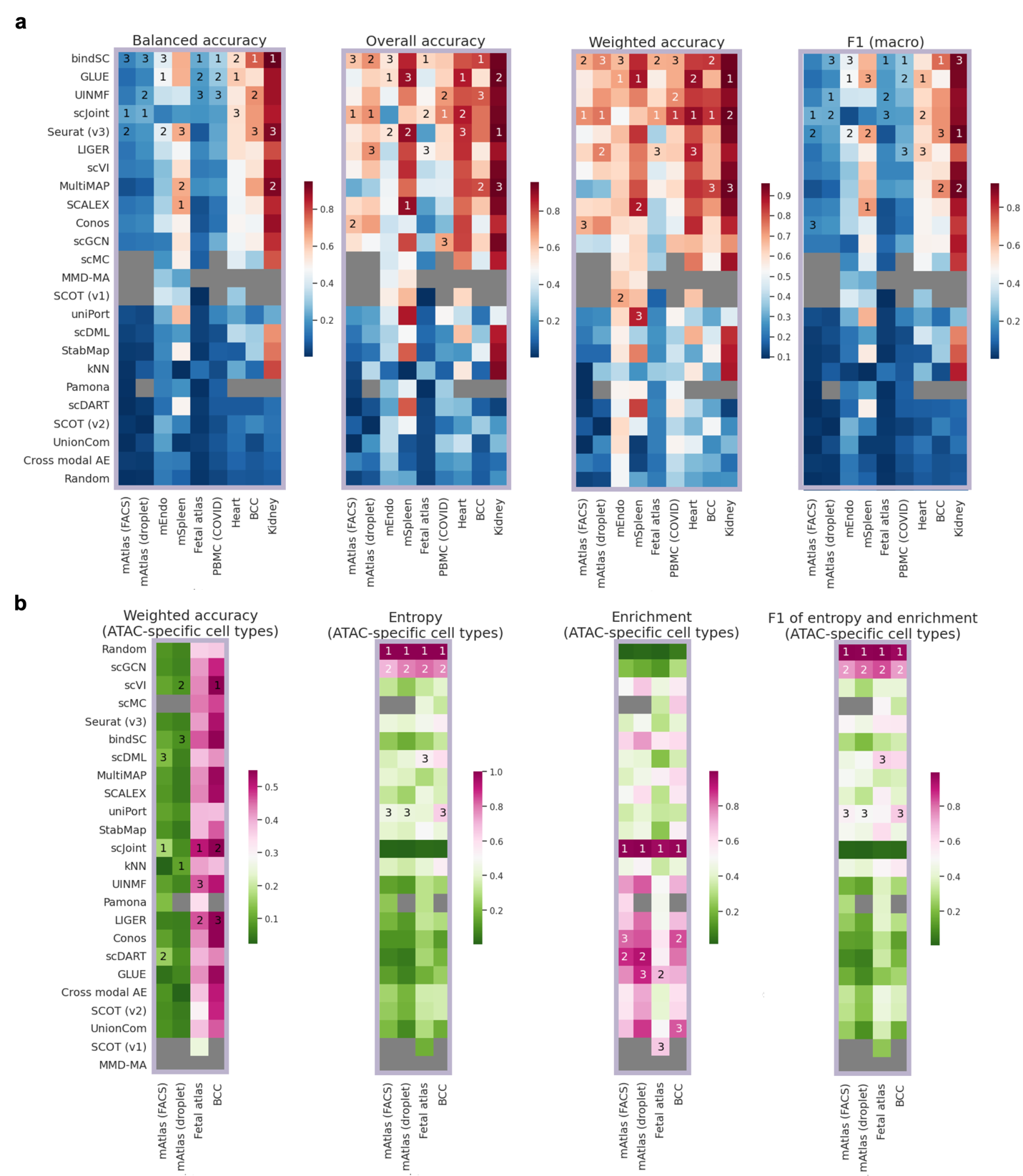

**Figure S3.** Detailed metrics on (a) common cells and (b) ATAC-specific cells for unpaired datasets without corresponding paired data. For each heatmap, methods (rows) are placed in the same order as that in Fig. 2b and methods ranked top three in each task are marked by Arabic numbers.

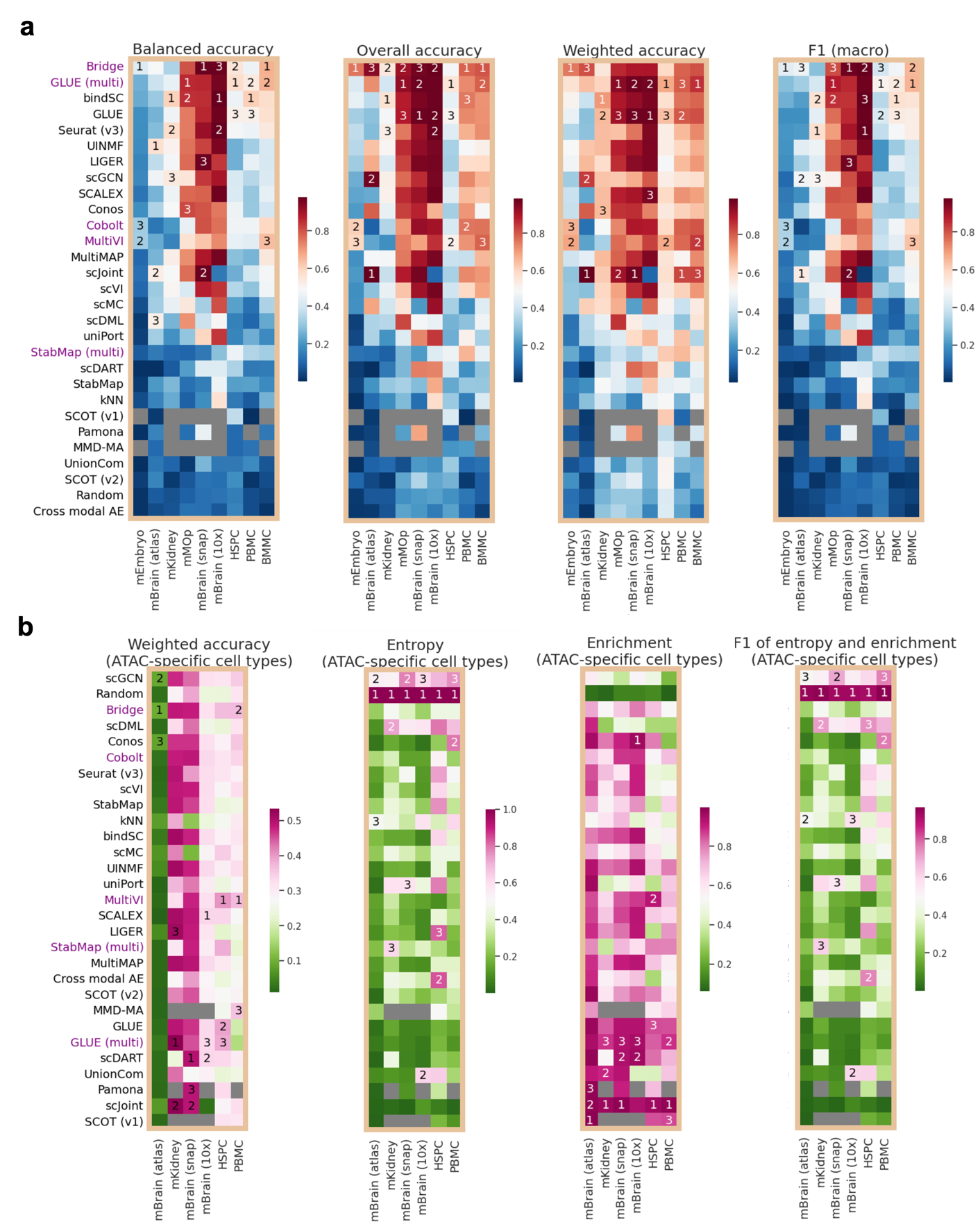

**Figure S4.** Detailed metrics on (a) common cells and (b) ATAC-specific cells for unpaired datasets with corresponding paired data. For each heatmap, methods (rows) are placed in the same order as that in Fig. 2c and methods ranked top three in each task are marked by Arabic numbers.

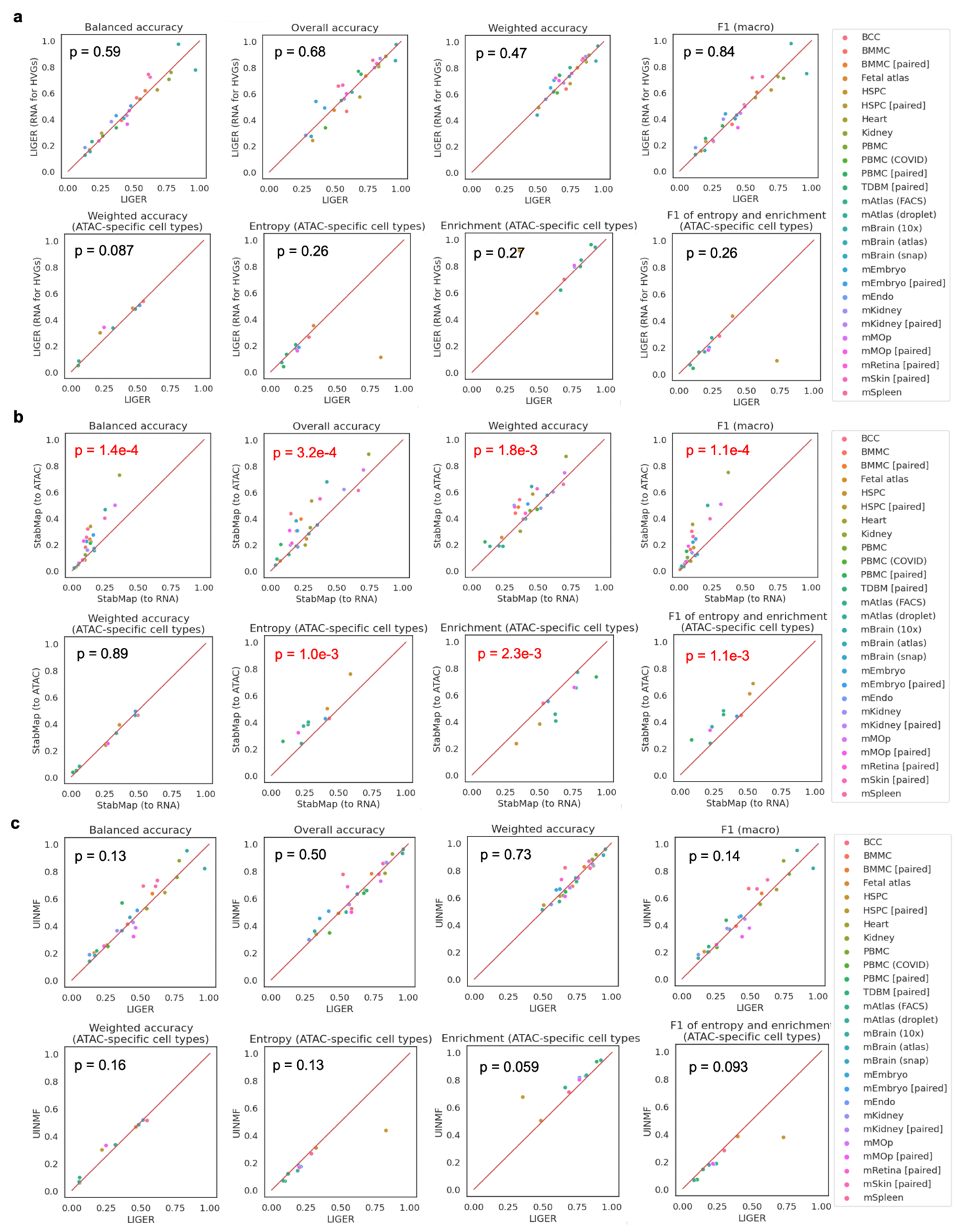

**Figure S5.** Some pair-wise comparisons between selected pairs of two methods or two versions of a method, including (a) LIGER and LIGER (use RNA for HVGs), (b) StabMap (to RNA) and StabMap (to ATAC), and (c) LIGER and UINMF.

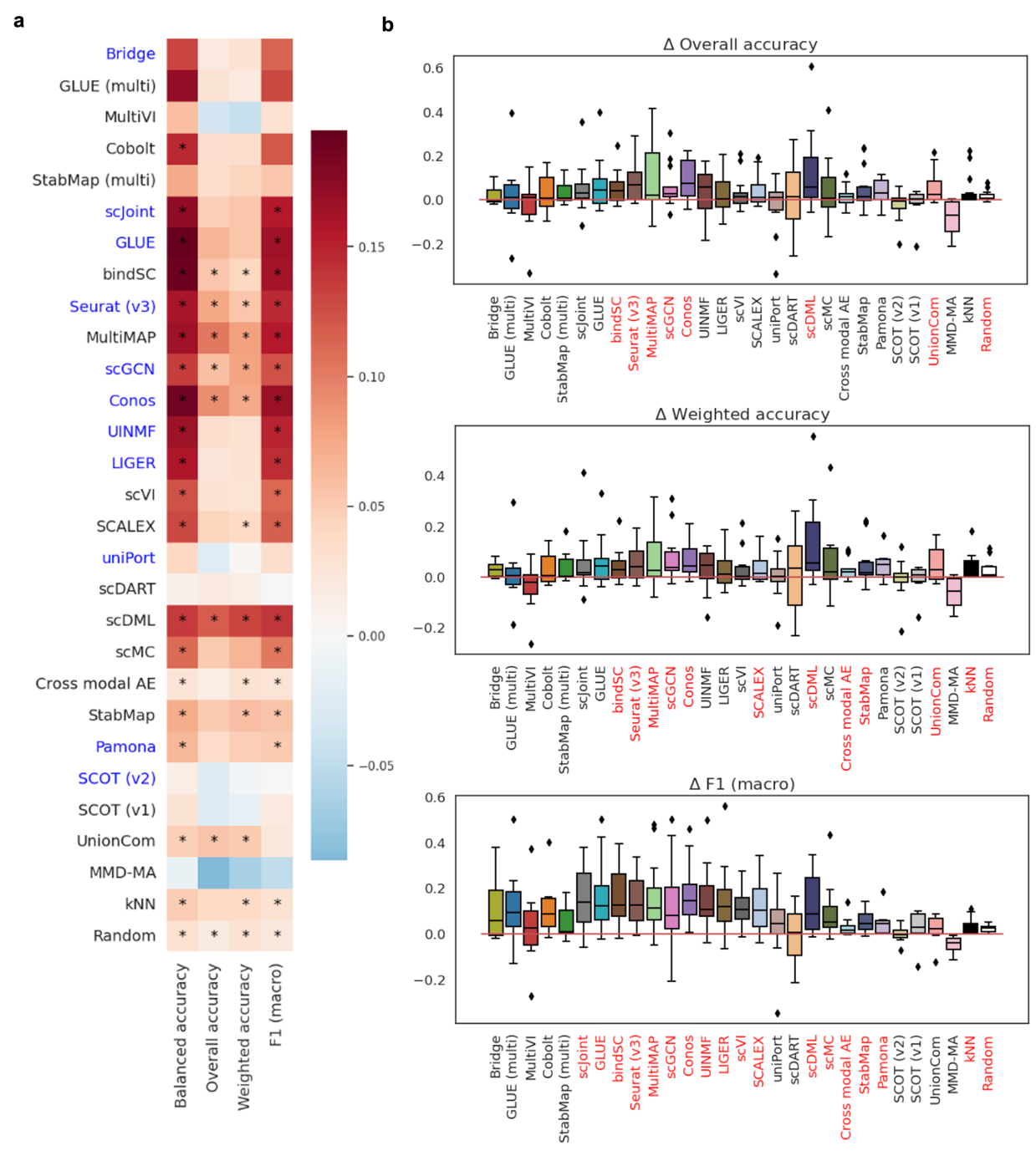

**Figure S6.** Supplementary information about the cell type imbalance experiment (common cell types - original). (a) The differences in accuracy metrics (columns) averaged across all tasks for the 27 methods (rows). Differences that achieved statistical significance are indicated by asterisks. Methods marked in blue are those that claim can deal with data imbalance. (b) Boxplots show the differences in overall accuracy and F1 (macro). Methods marked in red along the x-axes are those that achieved significant differences (p-value threshold: 0.05; statistical test: one-sample t-test).

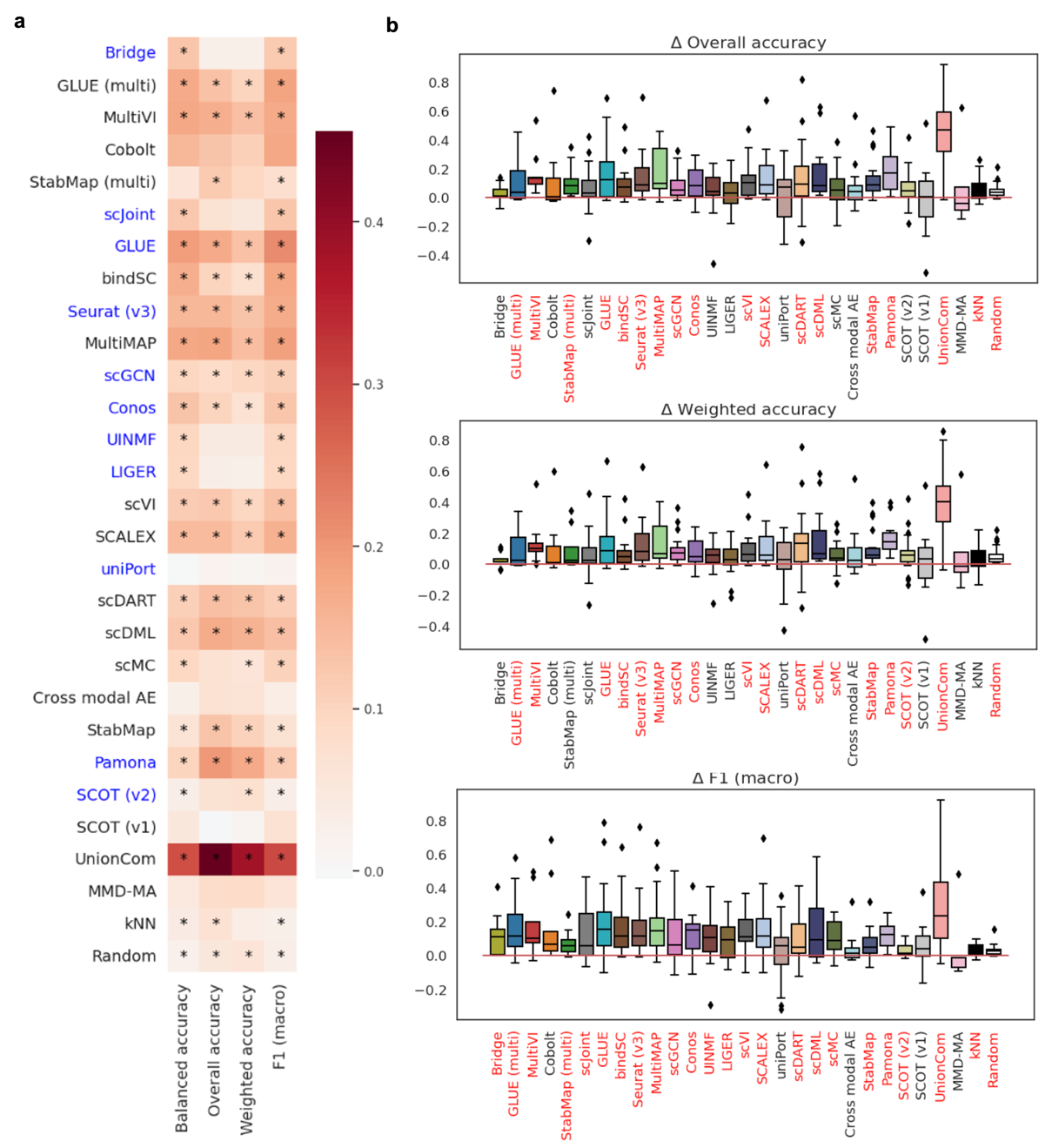

**Figure S7.** Supplementary information about the cell type imbalance experiment (balanced cell types - original). (a) The differences in accuracy metrics (columns) averaged across all tasks for the 27 methods (rows). Differences that achieved statistical significance are indicated by asterisks. Methods marked in blue are those that claim can deal with data imbalance. (b) Boxplots show the differences in overall accuracy and F1 (macro). Methods marked in red along the x-axes are those that achieved significant differences (p-value threshold: 0.05; statistical test: one-sample t-test).

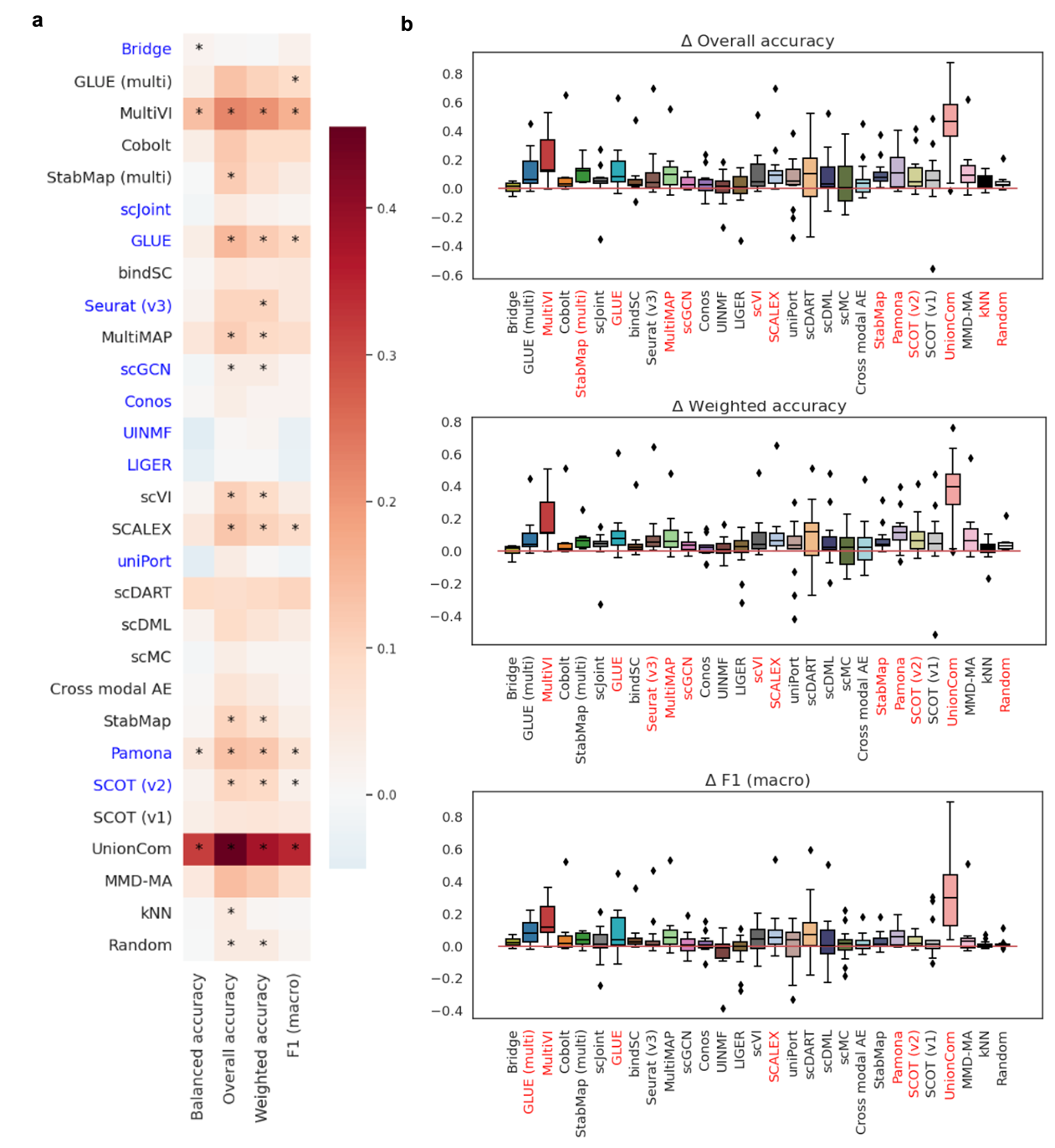

**Figure S8.** Supplementary information about the cell type imbalance experiment (balanced cell types - common cell types). (a) The differences in accuracy metrics (columns) averaged across all tasks for the 27 methods (rows). Differences that achieved statistical significance are indicated by asterisks. Methods marked in blue are those that claimed itself could deal with data imbalance. (b) Boxplots show the differences in overall accuracy and F1 (macro). Methods marked in red along the x-axes are those that achieved significant differences (p-value threshold: 0.05; statistical test: one-sample t-test).

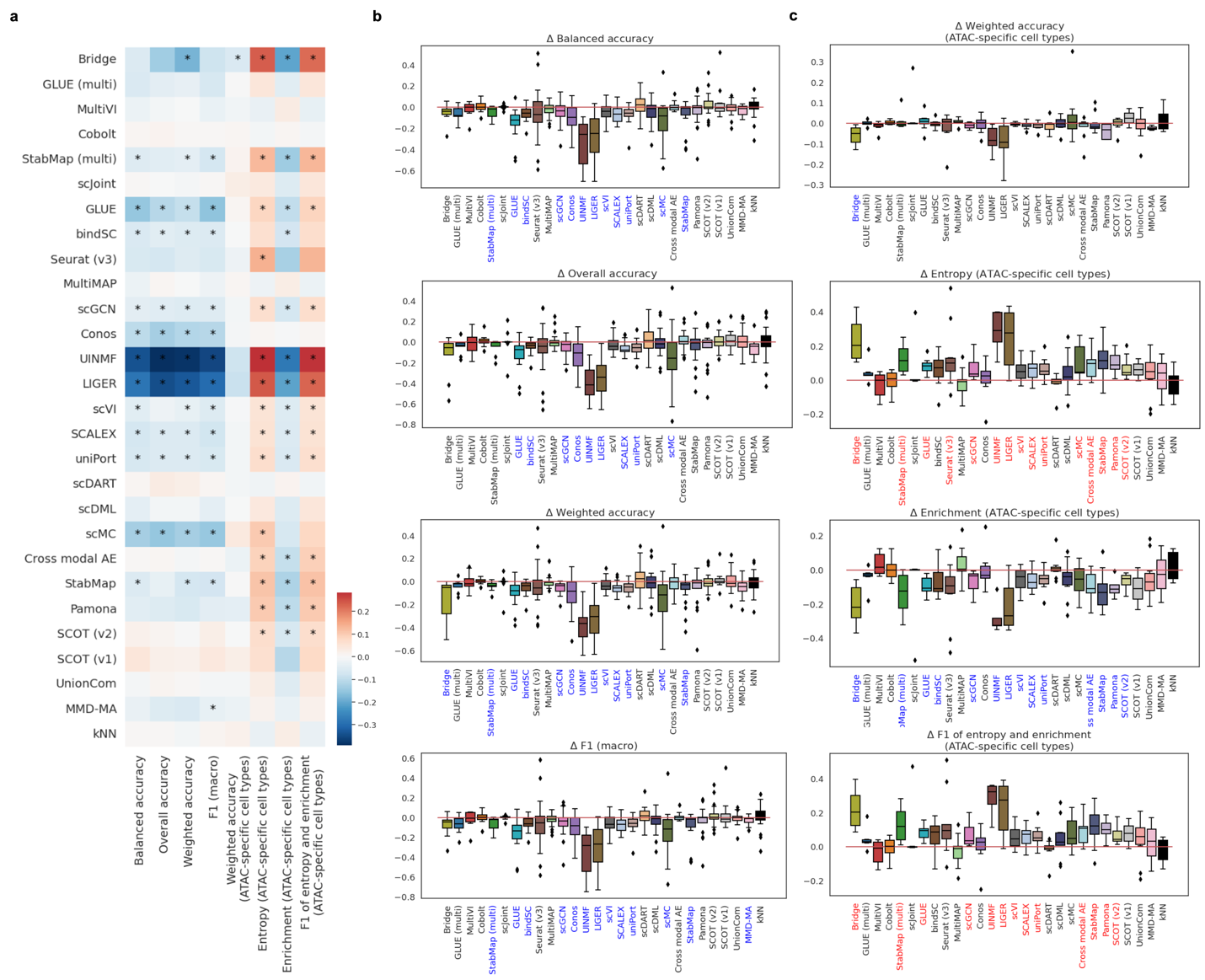

**Figure S9.** The impact of data binarization on all 27 methods evaluated by the differences between accuracy metrics calculated using results after and before data binarization. (a) The differences in accuracy metrics (columns) averaged across all tasks for the 27 methods (rows). Differences that achieved statistical significance are indicated by asterisks. (b) Boxplots show the differences in each of the four metrics calculated on common cells. (c) Boxplots show the differences in each of the four metrics calculated on ATAC-specific cells. In (b) and (c), methods marked in red and blue along the x-axes are methods that achieved significant positive and negative differences respectively (p-value threshold: 0.05; statistical test: one-sample t-test).

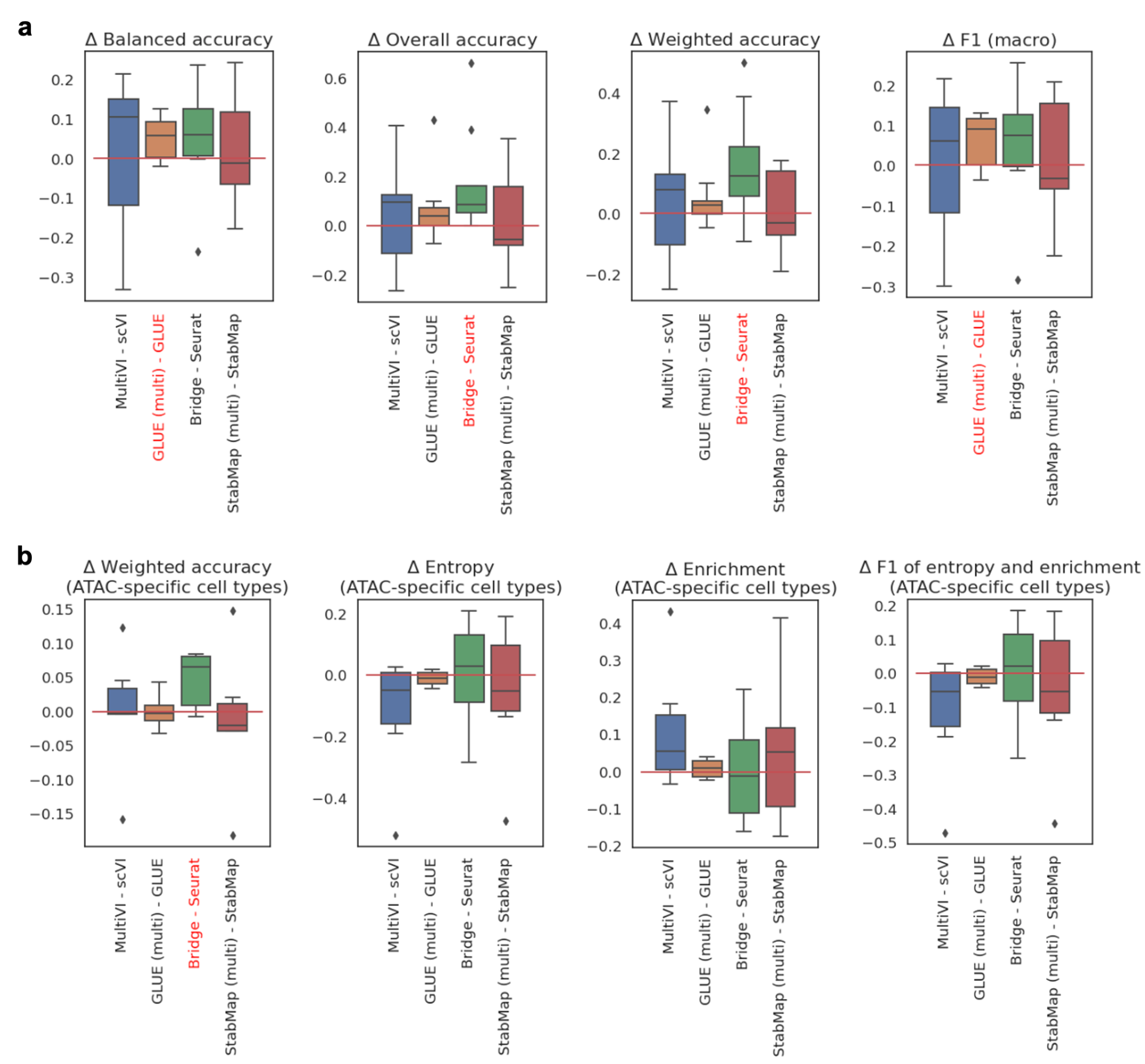

**Figure S10.** Supplementary information about the effect of using additional paired data as a 'bridge'. (a) Boxplots show the differences in the four metrics calculated on common cells. (b) Boxplots show the differences in the four metrics calculated on ATAC-specific cells. Methods marked in red along the x-axes are those that achieved significant differences (p-value threshold: 0.05; statistical test: one-sample t-test).

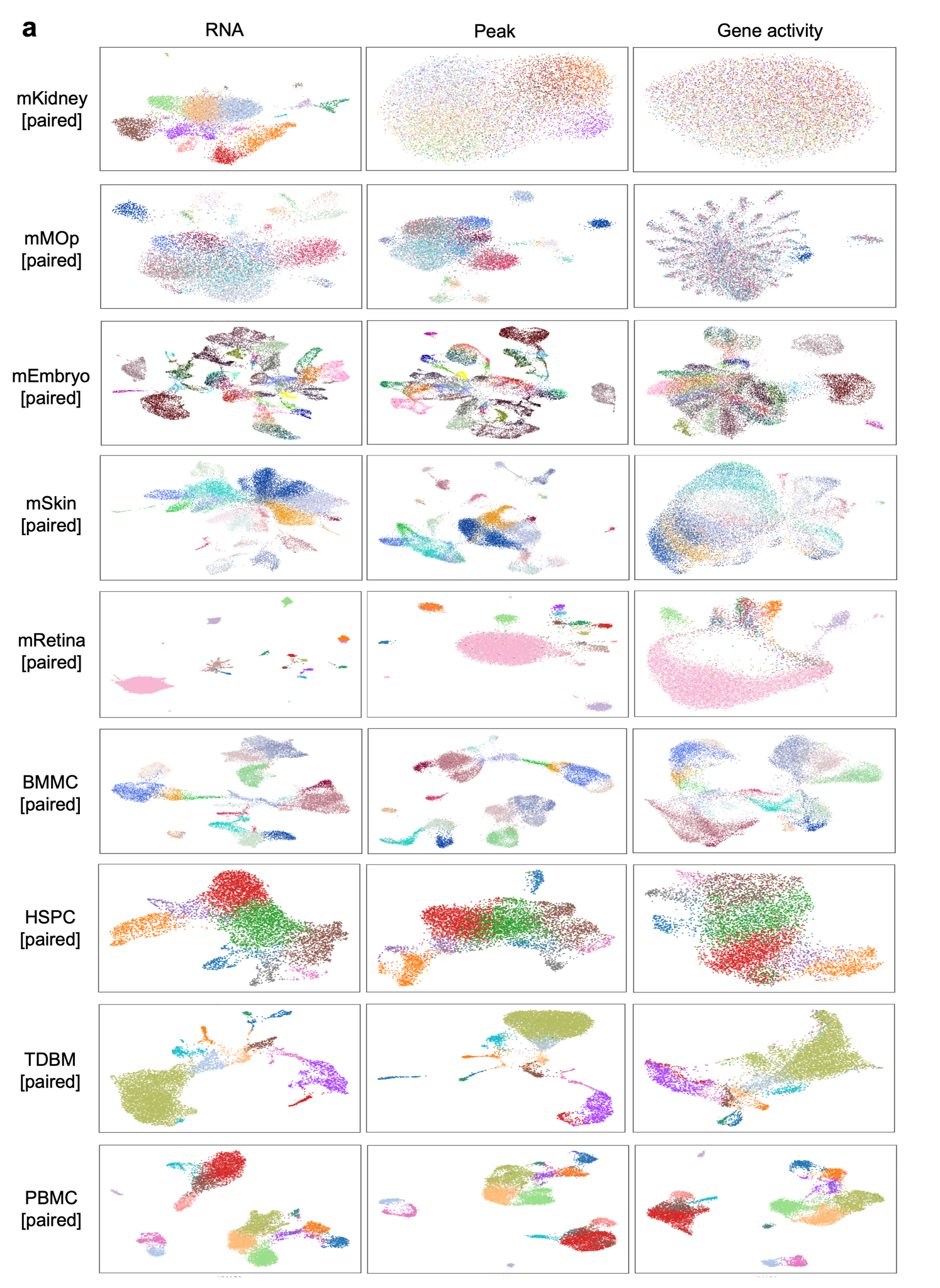

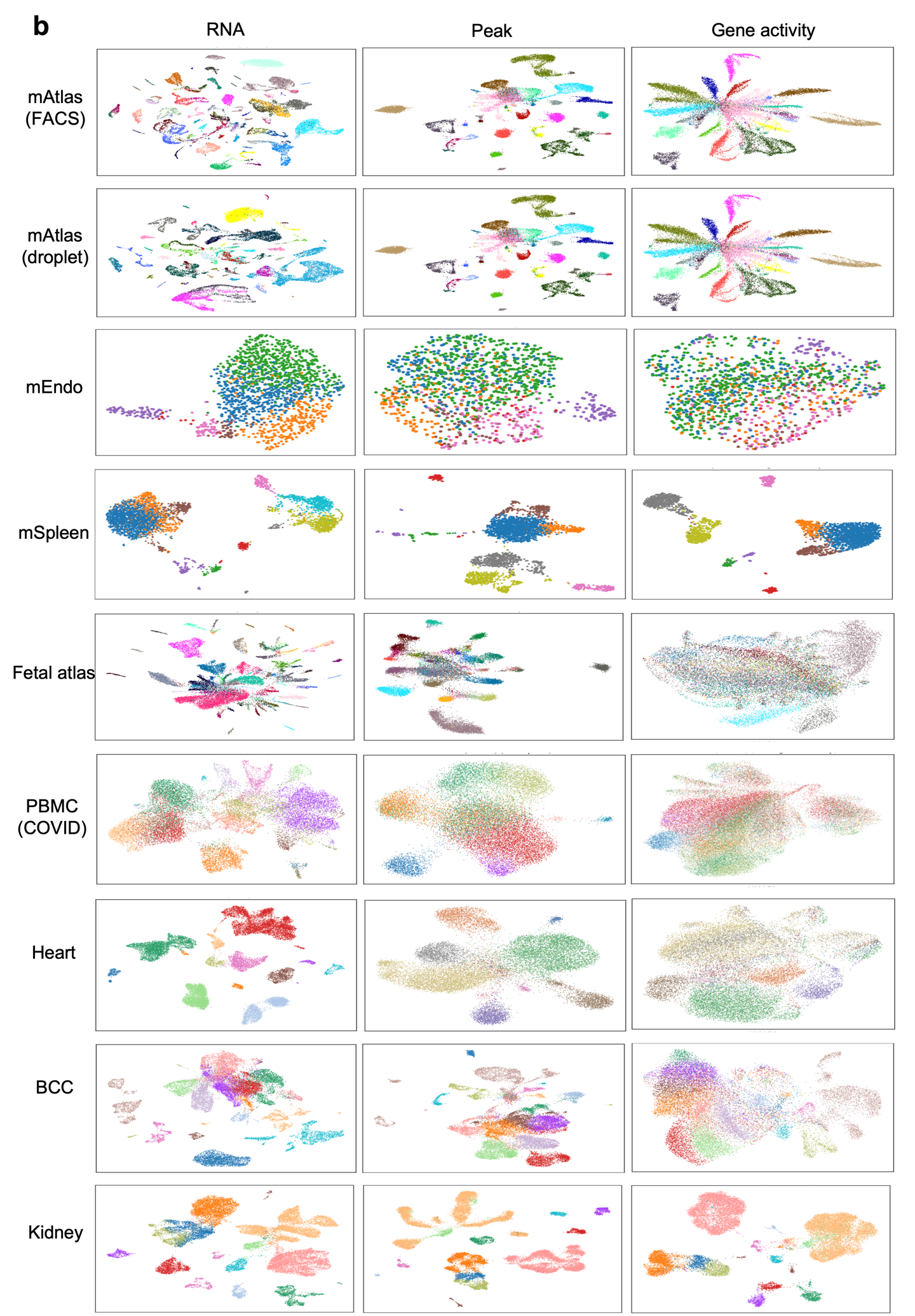

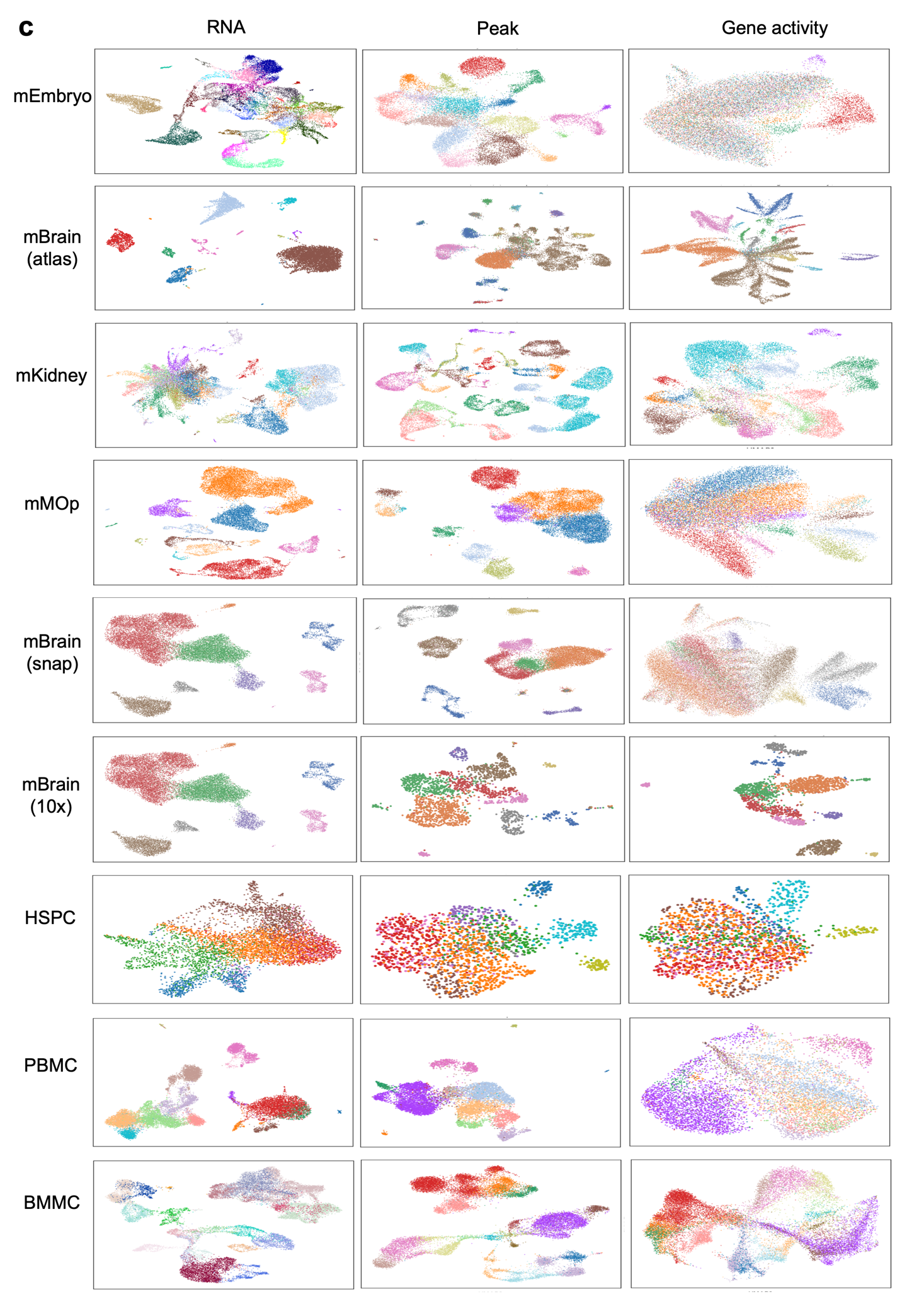

**Figure S11.** UMAPs plotted using RNA gene data (first column), ATAC peak data (second column) and ATAC gene activity data (third column). (a) Nine paired datasets. (b) Nine unpaired datasets that do not have corresponding paired data. (c) Nine unpaired datasets that have corresponding paired data.

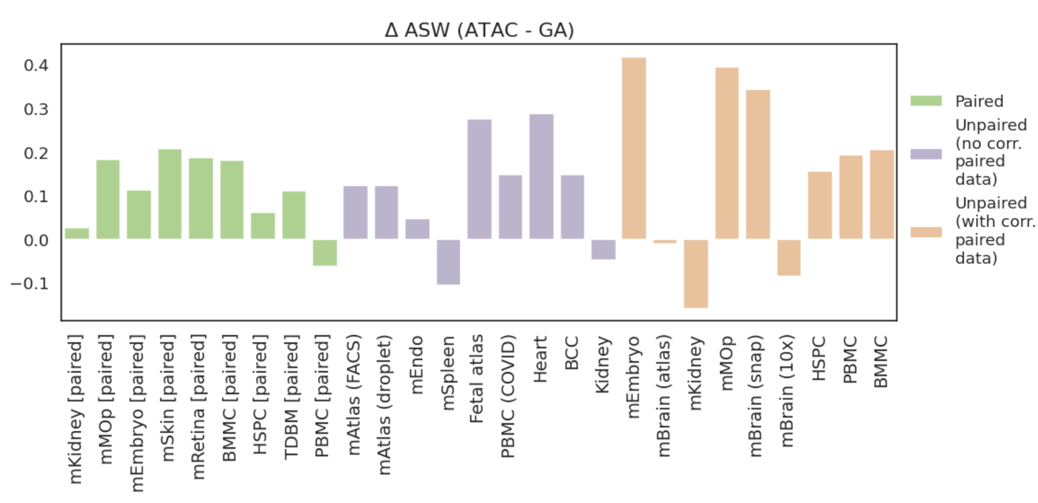

**Figure S12.** Supplementary figure for Fig. 4b. The absolute differences in ASW by subtracting ASW calculated using gene activity (GA) from ASW calculated using ATAC peak. The order of datasets is the same as that in Fig. 2.

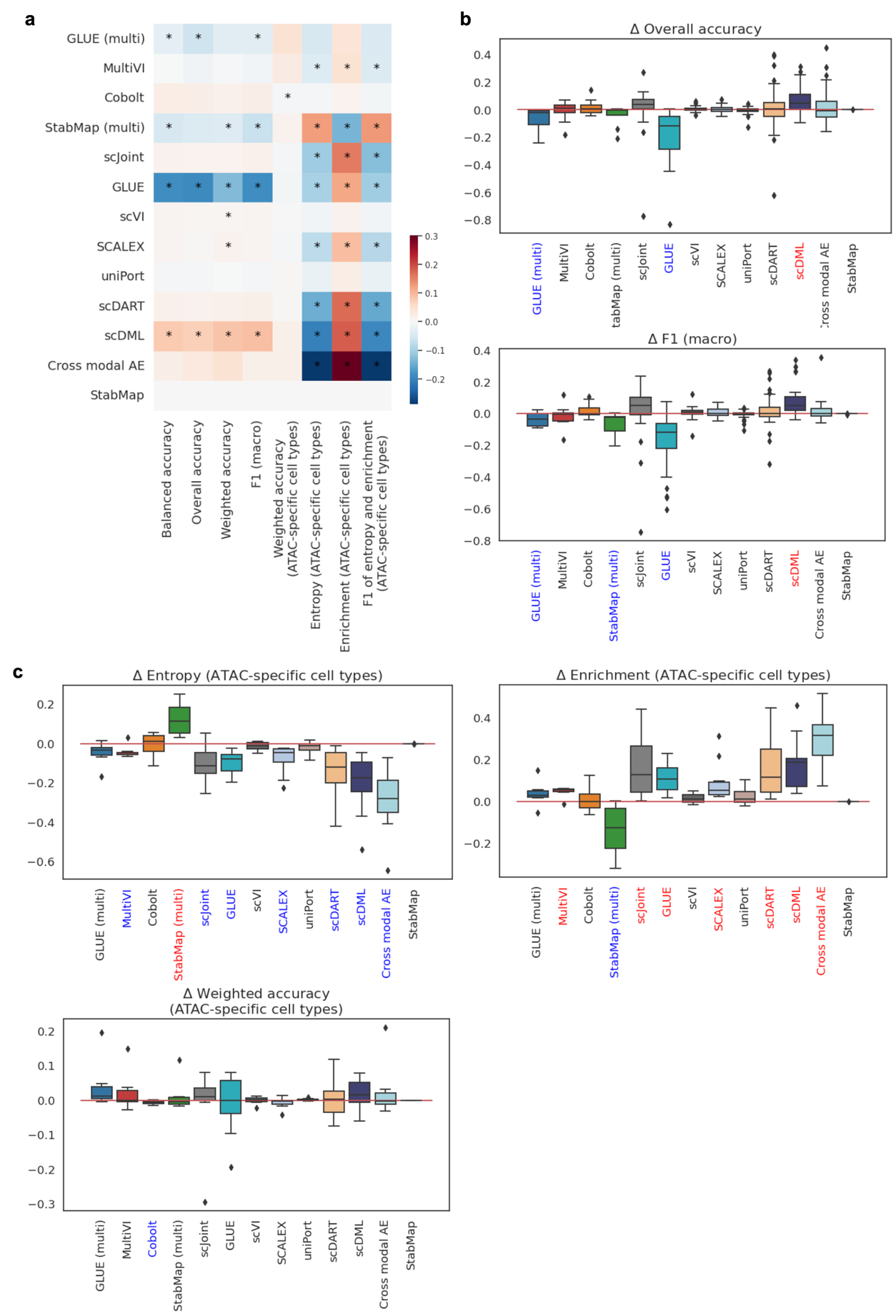

**Figure S13.** Supplementary information about the impact of introducing semi-supervised strategy to 13 selected methods on prediction accuracy. (a) The differences in accuracy metrics (columns) averaged across all tasks for the 13 methods (rows). Differences that achieved statistical significance are indicated by asterisks. (b) Boxplots show the differences in overall accuracy and F1 (macro) calculated on common cells. (c) Boxplots show the differences in entropy, enrichment and weighted accuracy calculated on ATAC-specific cells. In (b) and (c), methods marked in red and blue along the x-axes are those that achieved significant positive and negative differences respectively (p-value threshold: 0.05; statistical test: one-sample t-test).

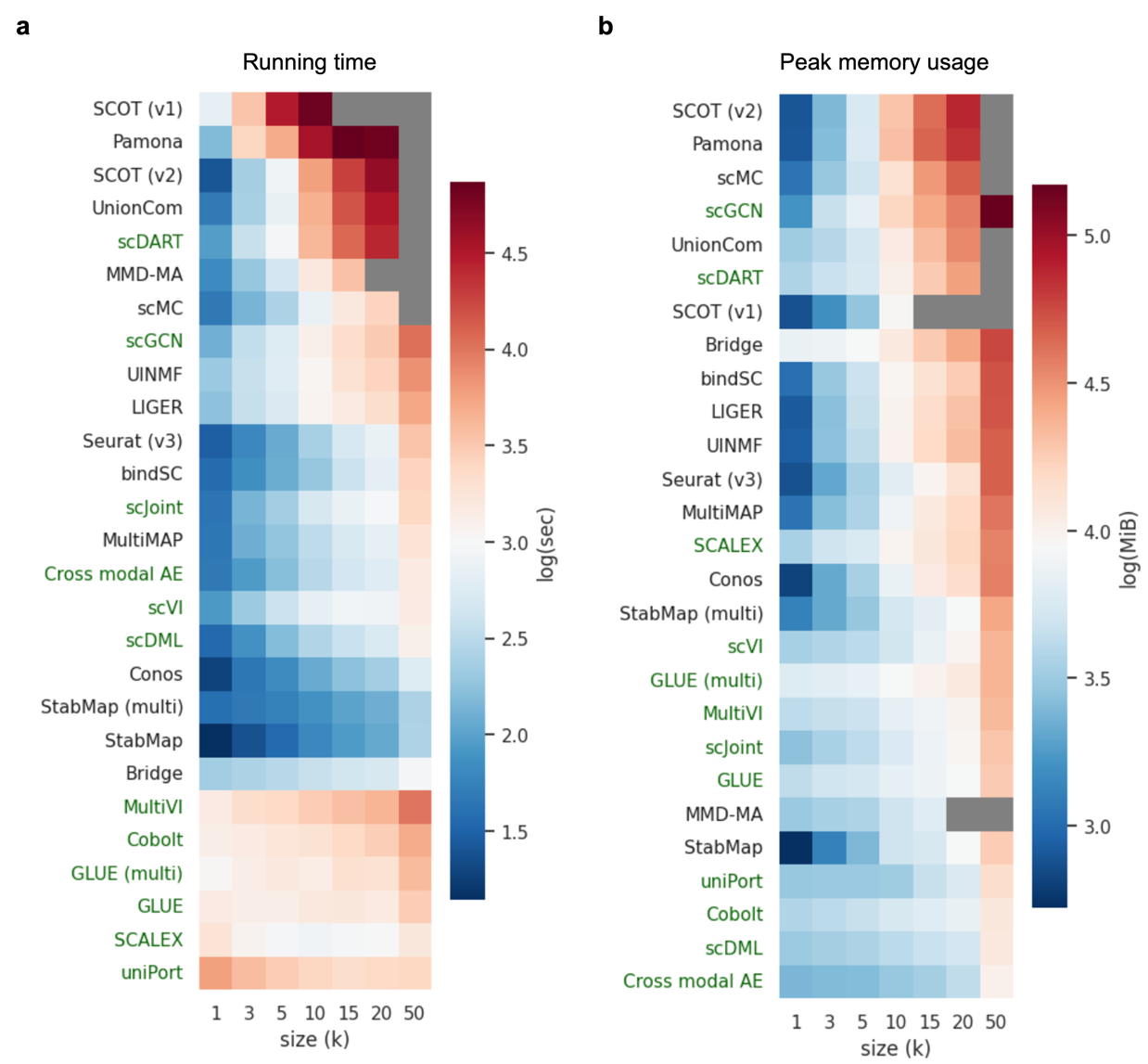

**Figure S14.** Supplementary information about computational efficiency. (a) Heatmap shows log-scaled running time in seconds. Grey elements are missing data because a method could not finish running in one day (SCOT (v1), Pamona, UnionCom and scDART), could not ran on GPU A100 (MMD-MA), consumed more memory than 200 GiB (SCOT (v2)), or failed (scMC). (b) Heatmap shows log-scaled peak memory usage in MiB. Grey elements are missing data because a method could not finish running in one day (SCOT (v1), Pamona, UnionCom and scDART), could not ran on GPU A100 (MMD-MA), consumed more memory than 200 GiB (SCOT (v2)), or failed (scMC).
